# An RNA binding regulatory cascade controls the switch from proliferation to differentiation in the *Drosophila* male germ cell lineage

**DOI:** 10.1101/2024.09.06.611673

**Authors:** Devon E. Harris, Jongmin J. Kim, Sarah R. Stern, Hannah M. Vicars, Neuza R. Matias, Lorenzo Gallicchio, Catherine C. Baker, Margaret T. Fuller

**Affiliations:** Department of Developmental Biology, Stanford University School of Medicine, Stanford, CA 94305, USA; Department of Chemical and Systems Biology, Stanford University School of Medicine, Stanford, CA 94305, USA; Department of Biomedical Sciences, Cornell University, Ithaca NY, 14853, USA; Department of Genetics, Stanford University School of Medicine, Stanford, CA 94305, USA

**Keywords:** differentiation, proliferation, spermatogenesis, RNA-binding proteins, adult stem cell lineage

## Abstract

The switch from precursor cell proliferation to onset of differentiation in adult stem cell lineages must be carefully regulated to produce sufficient progeny to maintain and repair tissues, yet prevent overproliferation that may enable oncogenesis. In the *Drosophila* male germ cell lineage, spermatogonia produced by germ line stem cells undergo a limited number of transit amplifying mitotic divisions before switching to the spermatocyte program that sets up meiosis and eventual spermatid differentiation. The number of transit amplifying divisions is set by accumulation of the *bag-of-marbles* (Bam) protein to a critical threshold. In *bam* mutants, spermatogonia proliferate through several extra rounds of mitosis then die without becoming spermatocytes. Here we show that a key role of Bam for the mitosis to differentiation switch is repressing expression of Held Out Wings (*how*), homolog of mammalian Quaking. Knockdown of *how* in germ cells was sufficient to allow spermatogonia mutant for *bam* or its partner *benign gonial cell neoplasm* (*bgcn*) to differentiate, while forced expression of nuclear-targeted How protein in spermatogonia wild-type for *bam* resulted in continued proliferation at the expense of differentiation. Our findings suggest that Bam targets *how* RNA for degradation by acting as an adapter to recruit the CCR4-NOT deadenylation complex via binding its subunit, Caf40. As How is itself an RNA binding protein with roles in RNA processing, our findings reveal that the switch from proliferation to meiosis and differentiation in the *Drosophila* male germ line adult stem cell lineage is regulated by a cascade of RNA-binding proteins.

## Introduction

In a common feature of the adult stem cell lineages that build and repair tissues throughout the body, relatively less differentiated precursor cells undergo a limited series of transit amplifying divisions, which serve to increase the number of differentiated progeny produced from a single adult stem cell division. Blood, skin, intestinal epithelia, and male germ cell lineages all employ transit amplifying (TA) divisions to produce large numbers of differentiated cells (1). In this context, the switch from proliferation to differentiation must be carefully regulated. Premature switching can lead to inadequate differentiated cell replacement, with defects in tissue maintenance or repair. Conversely, failure to switch resulting in overproliferation of precursor cells may increase danger of accumulating oncogenic mutations leading to cancer (2).

Genetic analysis of the *Drosophila* male germ line adult stem cell lineage has revealed key players in the switch from the mitotic program of transit amplifying spermatogonia to onset of meiosis and the spermatocyte gene expression program that sets up differentiation. Germ line stem cells at the testis apical tip divide with an oriented spindle (2) so that one daughter remains next to the apical hub and maintains stem cell identity, while the daughter displaced away from the hub becomes a gonialblast, founding a clone of transit amplifying spermatogonia (Figure 1A) (3). Because daughters of a single founding gonialblast divide in synchrony with incomplete cytokinesis and are enclosed by a pair of somatic cyst cells, counting the number of germ cells in each cyst can reveal the number of rounds of mitotic divisions that took place in the clone prior to the switch to the spermatocyte differentiation program. The number of transit amplifying mitotic divisions differs among *Drosophilid* species, so is under genetic control (4), with the number in wild-type *Drosophila melanogaster* almost always exactly four (Figure 1A).

**Figure 1.**
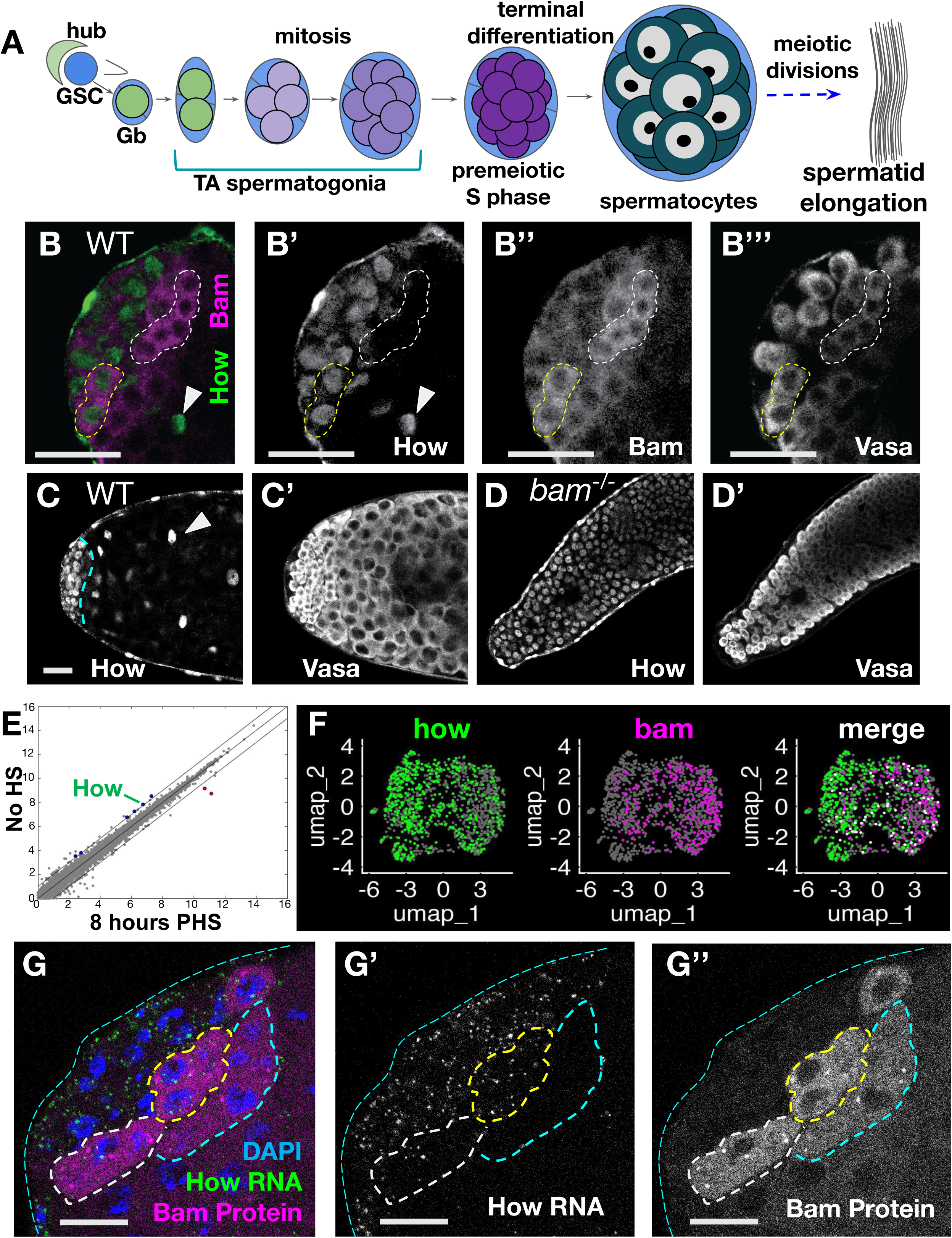
Nuclear How persists in *bam* mutant spermatogonia. (A) Diagram of *Drosophila* early male germ line development. (B-B’’’) Immunofluorescence images of apical tip of a wild type testis stained with anti-How, anti-Vasa, and anti-Bam. (B) merge of (green) How and (magenta) Bam. (B’-B’’’) Black and white single channels showing (B’) How, (B’’) Bam, and (B’’’) Vasa to mark germ cells. (C-D) Immunofluorescence images of (C) wild type and (D) *bam^1/^*^Δ*86*^ testis apical tips stained with anti-How and anti-Vasa. Dashed cyan line in (C) marks border between spermatogonia (left of line) and the spermatocyte region (right). Arrowheads in B and C: somatic cyst cell nuclei. (E) Scatter plot of RNA-sequencing data from *hs-Bam*; *bam^1/^*^Δ*86*^ comparing log_2_ transcript expression levels per gene for testes from flies not treated with heat shock (no HS) to testes from flies 8 hours post heat shock (PHS). (Blue dots) *how* is one of the 6 genes for which transcript levels decreased by over 2-fold by 8 hours post induction of Bam expression by heat shock in both the RNA-seq and separate microarray analysis (details in Figure S1). (Red dots) Transcripts from two genes increased in expression level in both the RNA-seq and microarray analysis. Diagonal lines mark 2-fold change. (F) UMAP visualization of single nuclear RNA sequencing data from the Fly Cell Atlas (14), after the nuclei in the two earliest male germ line clusters (Leiden resolution 6.0) were reclustered. (Green) nuclei scoring positive for *how*. (Magenta) nuclei scoring positive for *bam*. (White) nuclei positive for both *how* and *bam* transcripts. (G-G’’) Fluorescence images of testis apical tip showing (magenta) Bam-GFP protein and (green) *how* mRNA visualized by hybridization chain reaction (HCR) using probes to protein coding sequences of *how*. Dashed outlines: spermatogonial cysts positive for Bam protein either (yellow) with how RNA, (white) with fewer loci of how RNA signal, or (cyan) no how RNA signal detected. Scale bars: 25 μm in B-D and 12.5 μm in G.

Mutational analysis has identified three interacting proteins - Bag of marbles (Bam), Benign gonial cell neoplasm (Bgcn), and Tumorous testes (Tut), that are required for spermatogonia to stop proliferating and become spermatocytes (5–8). While Bgcn and Tut have RNA recognition motifs (8, 9), Bam does not contain any known RNA-binding domains. Bam appears to form a bridge in a ternary complex, with sequences in the Bam N-terminal third binding Tut and sequences in the Bam C-terminal third binding Bgcn (8). Bam protein expression detected by immunofluorescence staining showed the protein present in 4-cell cysts, higher in 8-cell cysts, and remaining high during premeiotic S phase, after which the protein quickly disappeared (4). Gene dosage experiments indicated that the number of transit amplifying divisions is set by the length of time it takes Bam protein to reach a critical threshold In flies with one mutant and one wild type allele of *bam*, spermatogonial cysts sometimes undergo one or occasionally more rounds of mitotic divisions before switching to spermatocyte differentiation. Conversely, in flies carrying a frameshift allele that deleted the Bam C-terminal PEST sequence, Bam protein accumulated more rapidly in transit amplifying spermatogonia and the cells often switched to spermatocyte state a round earlier, resulting in spermatocyte cysts with 8 germ cells rather than the normal 16 (4).

Bam protein has been shown to directly bind the Caf40 subunit of the CCR4-NOT deadenylation complex (CNOT9 in humans), suggesting that Bam may target RNAs for degradation or translational repression by recruiting the CCR4-NOT complex (10). While some targets of Bam in the male germ line are known, none of the genes identified so far appear to be key regulators in the switch from mitosis to meiosis downstream Bam. Previous work showed that Bgcn, Tut, and possibly Bam bind the 3’UTR of *mei-P26* and reduce expression of Mei-P26 protein. However, overexpression of *mei-P26* did not prevent the switch to spermatocytes, and decreasing *mei-P26* did not allow the switch to occur, suggesting that Mei-P26 is not the key target of Bam for the switch from mitosis to meiosis in males (8, 11).

Here we show that a key role of Bam and Bgcn in the switch from mitosis to meiosis in the *Drosophila* male germ line is to repress expression of the RNA-binding protein Held Out Wings (How), a homolog of mammalian Quaking, with roles in alternative RNA processing, export, and translation. How protein is normally downregulated in germ cells soon after Bam protein becomes expressed, but remained high in the TA spermatogonia that overproliferate in *bam* mutant males. Strikingly, knocking down *how* expression in *bam* mutant spermatogonia by RNAi allowed the germ cells to switch to spermatocyte state and differentiate. Reciprocally, forced expression of nuclear targeted How blocked differentiation of otherwise wild-type spermatogonia, even in the presence of Bam protein. Consistent with the model that Bam down regulates *how* RNA by recruiting Caf40 and the CCR4-NOT RNA degradation complex, knockdown of *Caf40* in early spermatogonia resulted in persistence of *how* RNA and overproliferation of TA cells, even in the presence of Bam protein. How is an RNA binding protein that regulates alternative RNA processing, export, and translation. Our findings reveal that an irreversible cell fate transition from mitosis to terminal differentiation in an adult stem cell lineage is controlled by a regulatory cascade of RNA-binding proteins.

## Results

### Downregulation of *how* is an early consequence of Bam action

Function of *bag-of-marbles* (*bam*) is required for proliferating spermatogonia to turn down expression of Held Out Wings (How). In testes wild-type for *bam*, immunofluorescence staining showed that How protein, present in the nucleus of early germ cells, is downregulated in mid-stage transit amplifying spermatogonia, soon after the Bam protein was first detected by immunofluorescence staining (Figure 1B), as previously documented by Monk *et al.* (2010) (12). Some mid-stage spermatogonia up regulating Bam protein still showed staining for How protein in their nuclei (Figure 1B, yellow dotted outline), while adjacent Bam positive cysts lacked detectable How protein (Figure 1B, white dotted outline). How protein continued to be detected in the nuclei of the somatic cyst cells that enclose spermatocyte cysts (Figure 1B, C, arrowheads), as well as in the testis sheath, but was below the level of detection in germ cells by early spermatocyte stages. In contrast, How protein persisted at high levels in the nuclei of the spermatogonia that continued to overproliferate in *bam* mutant males (Figure 1D).

When *bam* mutant spermatogonia were induced to differentiate in response to a burst of Bam expression under heat shock control, down regulation of *how* RNA was one of the earliest responses detected both by RNA-Seq and independent microarray analysis of whole testes Figures 1E; S1). In the heat shock Bam time course strategy developed by Kim *et al.*, (2017) (13), when *bam; hs-Bam* males were shifted as late pupae to 37°C for 30 minutes to induce expression of Bam then returned to 25°C, a wave of spermatogonia initiate differentiation, complete mitosis and a final S phase by 24 hours post heat shock (PHS), and differentiate into spermatocytes, with onset of expression of early spermatocyte-specific transcripts beginning by 24h PHS (Figure S1).

Analysis of PolyA+ RNAs expressed in whole testes at early time points in the heat shock-Bam differentiation time course showed that downregulation of *how* RNA was among the earliest changes detected. The level of *how* transcripts detected fell by > 2-fold by 8h PHS, long before the germ cells began to express spermatocyte-specific markers. *how* was one of six genes showing greater than 2-fold decrease in transcript level by 8h PHS by both RNA-sequencing and microarray analysis (Figure 1E and Figure S1A). In the gene expression comparisons in Figures 1E and S1, only those genes that met the following criteria were colored blue (for downregulated) or red (for upregulated): both RNA-Seq and independent microarray analysis of testes from *hs-Bam;bam* flies subjected to heat shock showed transcript levels changed >2 fold compared to the same genotype not subjected to heat shock, and testes from control *bam* mutant flies lacking the *hs-bam* did not show >2 fold change expression by microarray analysis for the same gene at the indicated times PHS compared to testes from *bam* flies not subjected to heat shock (Figure S1). Transcripts from additional genes became downregulated > 2-fold by later time points, with the number of genes with lowered transcripts growing from six at 8h PHS to 28 (16h), 83 (24h) and 114 by 32h PHS, by which time transcripts from a number of genes expressed specifically in spermatocytes began to be detected (Figure S1D,E).

Reclustering of snRNA-seq data from early germ cells generated by Li *et al.,* (14) showed that most early germ cell nuclei expressed either *how* RNA or *bam* RNA, while a small subset of the nuclei were positive for both (Figure 1F, white dots, merge). Fluorescence *in situ* hybridization (FISH) confirmed that levels of *how* RNA became down regulated as germ cells differentiate into spermatocytes (Figure S2B-E) and abruptly decreased in early germ cells soon after onset of Bam protein expression in wild-type testes (Figures 1G; S2C-E). In testes carrying a *Bam-GFP* transgene to allow visualization of Bam protein expression in mid to late transit amplifying spermatogonia, Hybridization Chain Reaction (HCR) FISH with probes recognizing the How protein coding sequence (Table 1) detected *how* transcripts in the nucleus and cytoplasm of cells near the testis tip, apical to the region where germ cells expressing Bam-GFP were detected (Figures 1G; S2C-E). *how in situ* signal was apparent in the cytoplasm of germ cells in some early spermatogonial cysts expressing Bam-GFP (yellow outline), while other Bam positive cysts showed little or no signal for *how* RNA (white and cyan outlines, respectively) (Figures 1G; S2C-E), suggesting that the level of *how* RNA in spermatogonia decreases rapidly once Bam protein becomes expressed. HCR FISH with a different probe set, directed against the 3’UTR associated with *how* mRNAs that encode nuclear targeted How(L) protein isoforms (Table 2) also detected *how* RNA signal at the testis apical tip that decreased with distance from the testis hub, like signal from the *how* protein coding region probe correlating with progression of transit amplifying spermatogonia to more mature stages (Figure S2B).

### *how* is a key target of Bam for the proliferation to differentiation switch

Knocking down expression of *how* in *bam* mutant TA spermatogonia by RNAi under control of *bam-Gal4* allowed *bam* mutant spermatogonia to differentiate into spermatocytes. Whereas squashed preparations of control testes wild-type for *bam* viewed by phase contrast microscopy showed abundant large spermatocytes and elongating spermatid bundles (Figure 2A), testes from males mutant for *bam* were considerably smaller, lacked spermatocytes and spermatids and contained large numbers of small germ cells that proliferate in cysts then eventually die (Figure 2B). However, if the *bam* mutant flies also carried a *UAS-how-RNAi* construct forcibly expressed in transit amplifying spermatogonia under control of *bamGal4*, the *bam*^-/-^ germ cells successfully differentiated into spermatocytes and elongating spermatids (Figures 2C; S3), indicating that the main requirement for Bam for the switch from mitosis to meiosis in males is reducing function of How. Similar rescue allowing differentiation of *bam* mutant germ cells to spermatocytes and elongating spermatids resulted from knockdown of expression of *how* by RNAi using a non overlapping shRNA construct from the TRiP collection (HMC03820) (Figure S3A-F). The *bam* mutant males in which *how* had been knocked down by *bamGal4* induced RNAi were fertile and produced viable offspring, similar to control males, although in both cases the number of progeny was quite low, possibly due to culturing the males at 29°C (Figure S3J).

**Figure 2.**
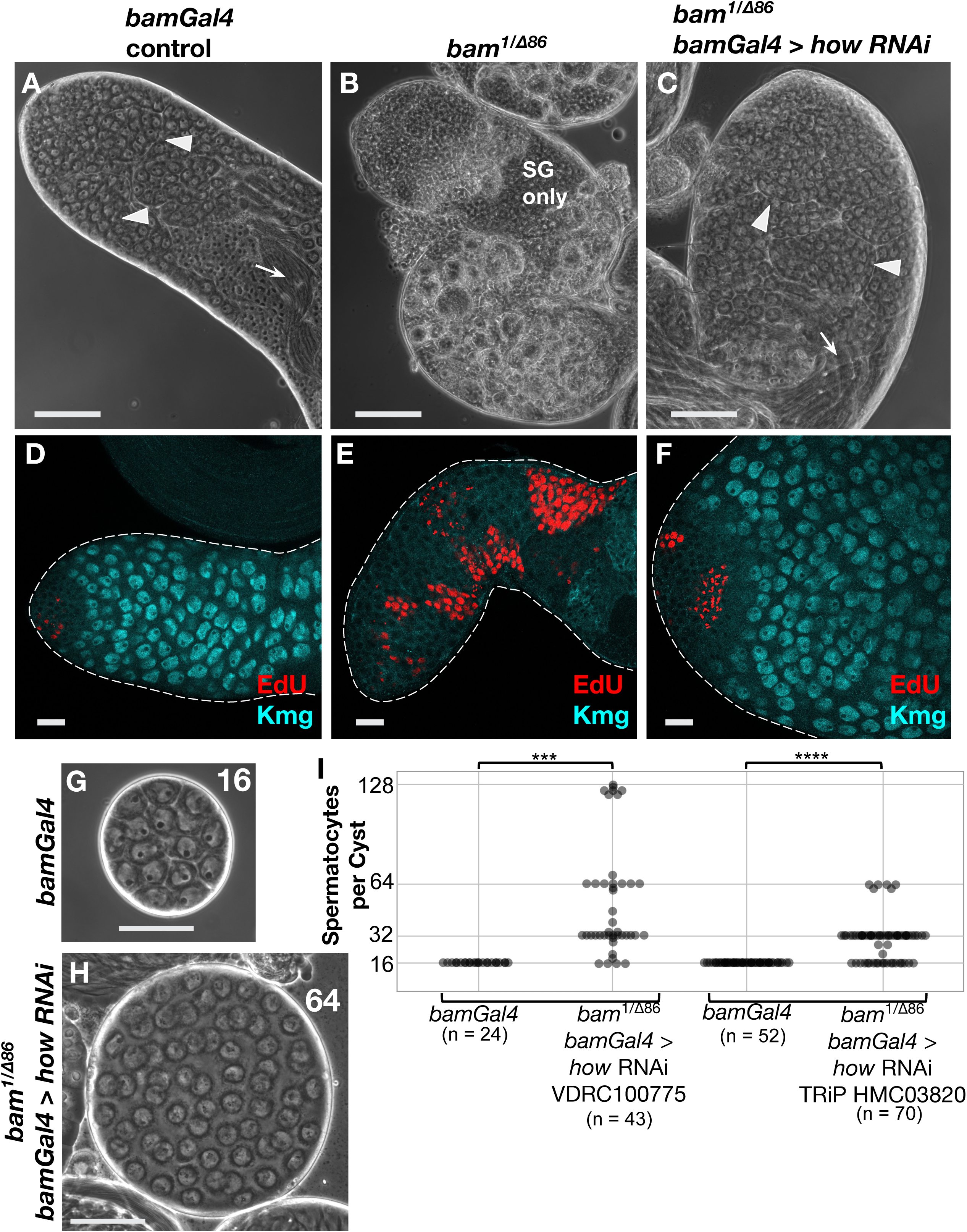
Knocking down *how* allowed spermatogonia lacking *bam* to differentiate into spermatocytes. (A-C) Phase contrast images of apical regions of testis in live squash preparations. (A) *bamGal4* expression driver only. (B) *bam^1/^*^Δ*86*^. (C) *bam^1/^*^Δ*86*^; *bamGal*4 > *how* RNAi (VDRC100775). All flies were raised under the same temperature shift regimen used for RNAi. Arrowheads: spermatocytes. Arrows: spermatid bundles. SG: spermatogonia. Scale Bars: 100 μm. (D-F) Immunofluorescence images of testis apical tips stained for (red) EdU to mark nuclei in S phase and (blue) Kmg to mark spermatocyte nuclei. (D) *bamGal4* driver only control. (E) *bam^1/^*^Δ*86*^. (F) *bam^1/^*^Δ*86*^; *bamGal4* > *how* RNA. Scale Bars: 25 μm. (G, H) Phase contrast images of intact spermatocyte cysts marked with the number of spermatocytes in the cyst. (G) *bamGal4* driver only. (H) *bam^1/^*^Δ*86*^; *bamGal4* > *how* RNAi. Scale Bars: 50 μm. (I) Number of spermatocytes per cyst from: (Left side) *bamGal4* sibling controls (N=24 cysts) or *bam^1/^*^Δ*86*^; *bamGal4* > *how RNAi* VDRC100775 (N=43 cysts). (***) p-value = 1.716e-10, based on a Wilcoxon rank-sum test with continuity correction. (Right side) *bamGal4* control (N=52 cysts) or *bam^1/^*^Δ*86*^; *bamGal4* > *how RNAi* TRiP HMC03820 from Bloomington Stock #55665 (N=70 cysts). (****) p-value = 1.033e-14 based on a Wilcoxon rank-sum test with continuity correction.

Testes from males mutant for *bgcn*, a binding partner of *bam*, had the same phenotype as *bam* mutants, overproliferation of transit amplifying spermatogonia that maintained expression of nuclear How protein (Figure S4A), and eventual germ cell death, with no cells at later stages of development (Figure S4B). Knockdown of *how* function in mid-to-late transit amplifying spermatogonia by RNAi under control of *bamGal4* also restored ability of *bgcn* mutant spermatogonia to differentiate into spermatocytes and elongating spermatids (Figure S4C).

Brief incubation of testes in EdU to label cells in S phase and immunofluorescence staining for the spermatocyte marker Kmg confirmed that knockdown of *how* by RNAi in mid-to-late spermatogonia under control of *bamGal4* restored the ability of *bam* mutant spermatogonia to switch to the spermatocyte state. Testes from control or RNAi knockdown flies were incubated in EdU for 5 minutes to label nuclei undergoing DNA replication, then immediately fixed and processed for immunofluorescence staining and imaging. Control testes from flies carrying the *bamGal4* transgene and raised under the same temperature regimen used for RNAi knockdowns showed a few small clusters of EdU-positive nuclei located near the testis apical tip, marking cysts of spermatogonia undergoing S phase in synchrony. As expected, no EdU positive nuclei were identified further from the tip, where germ cells showed expression of the spermatocyte specific marker Kmg (Figure 2D). In contrast, testes from flies mutant for *bam* raised under the RNAi knockdown temperature shift regime showed many more EdU-positive cysts per testis, several with more than 16 EdU-positive nuclei, indicating spermatogonial overproliferation, and no spermatocytes expressing Kmg (Figure 2E). Testes from *bam* mutant males in which expression of *how* was knocked down in late spermatogonia and early spermatocytes by RNAi under control of *bamGal4* showed many fewer EdU-positive cysts per testis, again confined to near the testis apical tip, and abundant Kmg-positive spermatocytes (Figure 2F). Similar results were observed when expression of How was knocked down by a different RNAi construct expressed under control of *bamGal4* (Figure S3D-F).

Notably, testes from *bam*^-/-^; *bamGal4;UAS-How RNAi* males raised under knockdown conditions were unusually wide, with many early germ cells (Figures 2C, F; S3I). The increased width of *bam*^-/-^; *bamGal4;UAS-How RNAi* testes was likely because downregulation of *how* due to RNAi under control of *bamGal4* may occur later than in 4-8 cell cysts as in wild type, because of the time it takes for the *bamGal4* to be expressed, activate transcription of the RNAi construct, then for the RNAi to act upon *how* transcripts. Such delay would allow the *bam* mutant spermatogonia to go through additional rounds of mitotic transit amplifying divisions before downregulation of How. Consistent with this, spilling out individual cysts revealed that control testes raised under the knockdown temperature regimen almost always had 16 spermatocytes per cyst. In contrast, *bam^-/-^*; *bamGal4;UAS-How RNAi* testes normally had 32, 64, and sometimes more spermatocytes per cyst, indicating five, six, or more rounds of transit amplifying divisions prior to the switch to spermatocyte state, rather than the normal four (Figure 2G-I).

Conversely, forced expression of nuclear-targeted How but not cytoplasmic How in mid-stage spermatogonia was sufficient to largely block differentiation of otherwise wild-type spermatogonia into spermatocytes. The *how* locus encodes several transcript and protein isoforms (Figure S2A) (15). Some transcripts encode long forms of How protein that have a C-terminal nuclear localization signal (How(L)). Others encode shorter forms of the protein that lack the nuclear localization signal (How(S)) and are cytoplasmic.

Flies bearing a transgene with UAS sequences controlling inducible expression of How(L) protein (isoform A in Figure S2A) cloned in frame with a C-terminal HA epitope tag followed by terminator sequences from SV40 (*UAS-How(L)HA-SV40*) were crossed to flies bearing *bamGal4* to drive expression in mid-to-late transit amplifying spermatogonia. The mated flies were cultured at 25°C for three days then adults were removed and the progeny shifted to and maintained at 29°C to boost expression from the *UAS-How(L)HA-SV40* transgene. Under these conditions, testes from the newly eclosed *bamGal4; UAS-How(L)-HA-SV4* progeny had extensive apical regions filled with small germ cells (Figure 3B red bracket, compare to A), followed by areas with dying cells, as in *bam* mutant testes. Some testes also had a few cysts containing spermatocytes or post-meiotic spermatids, usually located far down the testis away from the apical tip, distal to the region with cell death.

**Figure 3.**
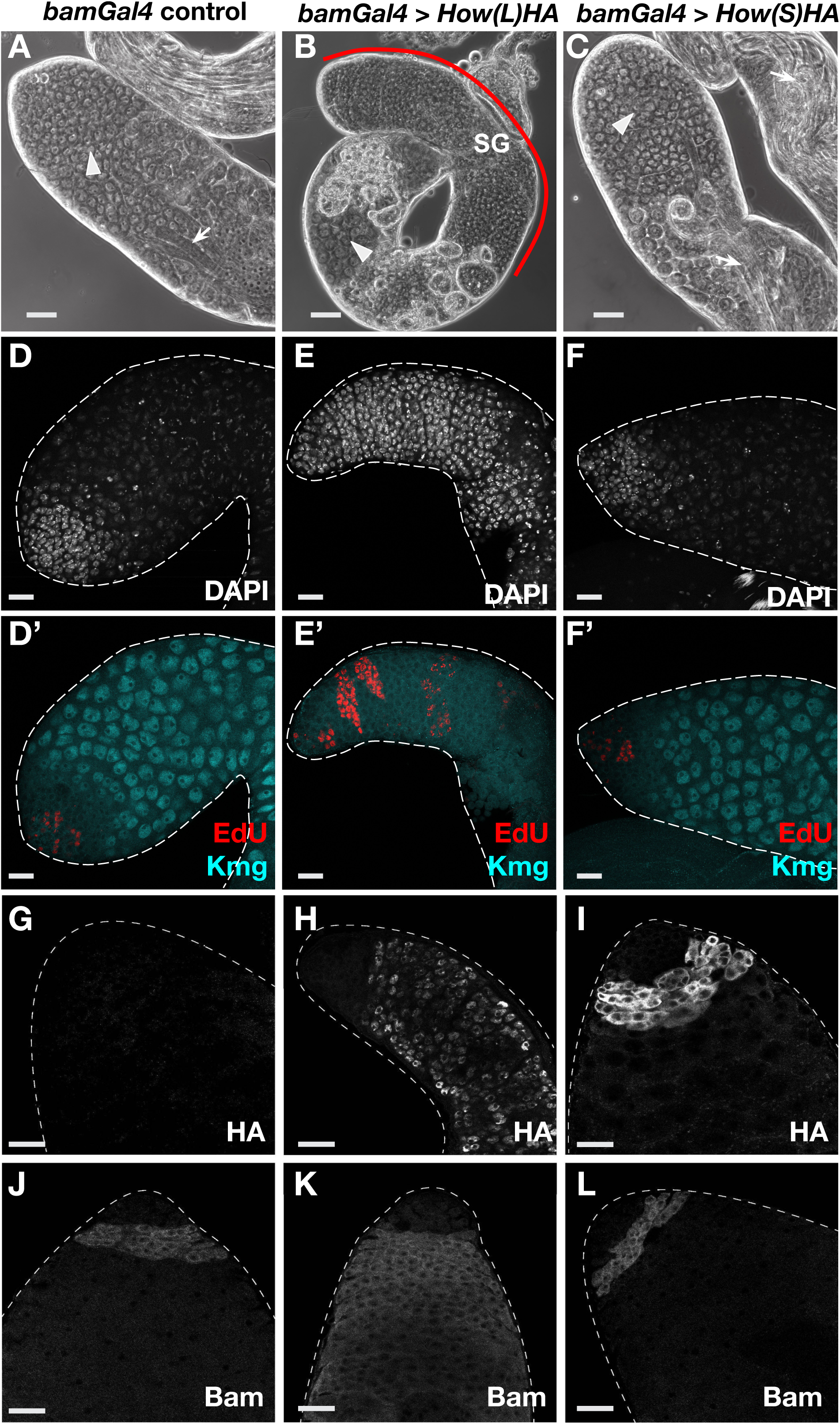
Overexpression of How(L) but not How(S) blocked the transition from spermatogonia to spermatocytes. (A-C) Phase contrast images of testis apical tips from (A) *bamGal4* driver only. (B) *bamGal4* > *UAS-How(L)HA-SV40*. (C) *bamGal4* > *UAS-How(S)HA-SV40*. SG: spermatogonia. Arrowheads: spermatocytes. Arrows: elongating spermatids. Red line in B marks region with overproliferating spermatogonia. Scale bars: 50 μm. (D-F) Immunofluorescence images of apical tips of testes stained with (top row) DAPI to mark nuclei and (bottom row) EdU to mark nuclei in S phase and anti-Kmg to mark spermatocyte nuclei. (D) *bamGal4* driver only. (E) *bamGal4* > *UAS-How(L)HA-SV40*. (F) *bamGal4* > *UAS-How(S)HA-SV40*. Scale Bars: 25 μm. (G-I) Immunofluorescence images of testis apical tips stained with anti-HA to detect the forcibly expressed How(L) and How(S). (G) *bamGal4* driver alone control. (H) *bamGal4*, *UAS-How(L)HA-SV40* showing nuclear localization of How(L). Note persistence of How(L)HA expression in the overproliferating spermatogonia. (I) *bamGal4* > *UAS-How(S)HA-SV40* showing cytoplasmic localization of How(S). Note shut off of How(S)HA expression in differentiating spermatocytes. (J-L) Immunofluorescence images of testis apical tips stained with anti-Bam. (J) *bamGal4* control. (K) *bamGal4* > *UAS-How(L)HA-SV40*. (L) *bamGal4* > *UAS-How(S)HA-SV40*. Scale bars: 25 μm.

Immunofluorescence staining after a short pulse of EdU revealed that most (∼80%) of the *bamGal4; UAS-How(L)HA SV40* testes had a much larger than normal number of cysts with germ cells undergoing synchronous DNA replication per testis than controls (Figure 3E’). The germ cell cysts subjected to forced expression of How(L) often had more than 16 nuclei undergoing S phase, indicating overproliferation. In addition, the nuclei remained small and stained brightly with DAPI, unlike in spermatocytes, and lacked expression of the spermatocyte marker Kmg (Figure 3E’). Immunofluorescence staining with anti-HA confirmed that How(L)-HA was localized to germ cell nuclei and that the *bamGal4* driver did not force expression of *How(L)HA* in germ line stem cells and early spermatogonia (Figure 3H). Notably, immunofluorescence staining with anti-Bam showed that Bam protein was expressed in the spermatogonia that overproliferated when *How(L)HA-SV40* was forcibly expressed, indicating that the failure of spermatogonia to differentiate was not due to repression of Bam by nuclear targeted How(L) (Figure 3K).

In contrast, testes from males expressing *UAS-How(S)HA-SV40* (isoform B in Figure S2A, which lacks the NLS) under control of *bamGal4* raised under the same conditions did not show massive overproliferation of spermatogonia, but instead had a modest population of small germ cells at the testis apical tip followed by plentiful differentiating spermatocytes, visible as large cells with large nuclei in unfixed squashed preparations viewed by phase contrast microscopy (Figure 3C). Immunofluorescence staining confirmed that the switch to spermatocyte state occurred after a limited number of mitotic divisions when *How(S)HA* was forcibly expressed in mid-to-late spermatogonia. Testes from *bamGal4; UAS-How(S)HA-SV40* males subjected to a brief incubation in EdU showed only a small number of EDU positive cysts, all of which were close to the apical tip. The testes also had many spermatocytes, marked by large nuclei positive for Kmg protein, starting from within a few cell diameters of the testis apical tip (Figure 3F’). Immunofluorescence staining with anti-HA confirmed that How(S)HA was cytoplasmic (Figure 3I). As expected for the *bamGal4* driver, anti-HA was not detected in very early germ cells at the testis tip. Together these data indicate that it is nuclear rather than cytoplasmic forms of How protein that block ability of spermatogonia to differentiate into spermatocytes.

Addition of 3’UTR sequences from *how(L)* mRNA reduced the effect of forced expression of *How(L)HA* on ability of spermatogonia to stop proliferating and differentiate into spermatocytes. *How(L)* mRNA isoforms have overlapping 1566nt - 2424nt 3’UTRs, which do not share sequences with the much shorter 3’UTRs of mRNA isoforms that encode How(S) proteins (15) (see Figure S2A). The 2424nt How(L) 3’UTR from *how* mRNA isoform A (Figure S2A) was cloned into the *UAS-How(L)HA-SV40* overexpression construct between the HA tag and the SV40 terminator to generate *UAS-How(L)HA-How(L)3’UTR-SV40* (Figure 4, top diagram) and introduced into flies as a transgene (Materials and Methods).

**Figure 4.**
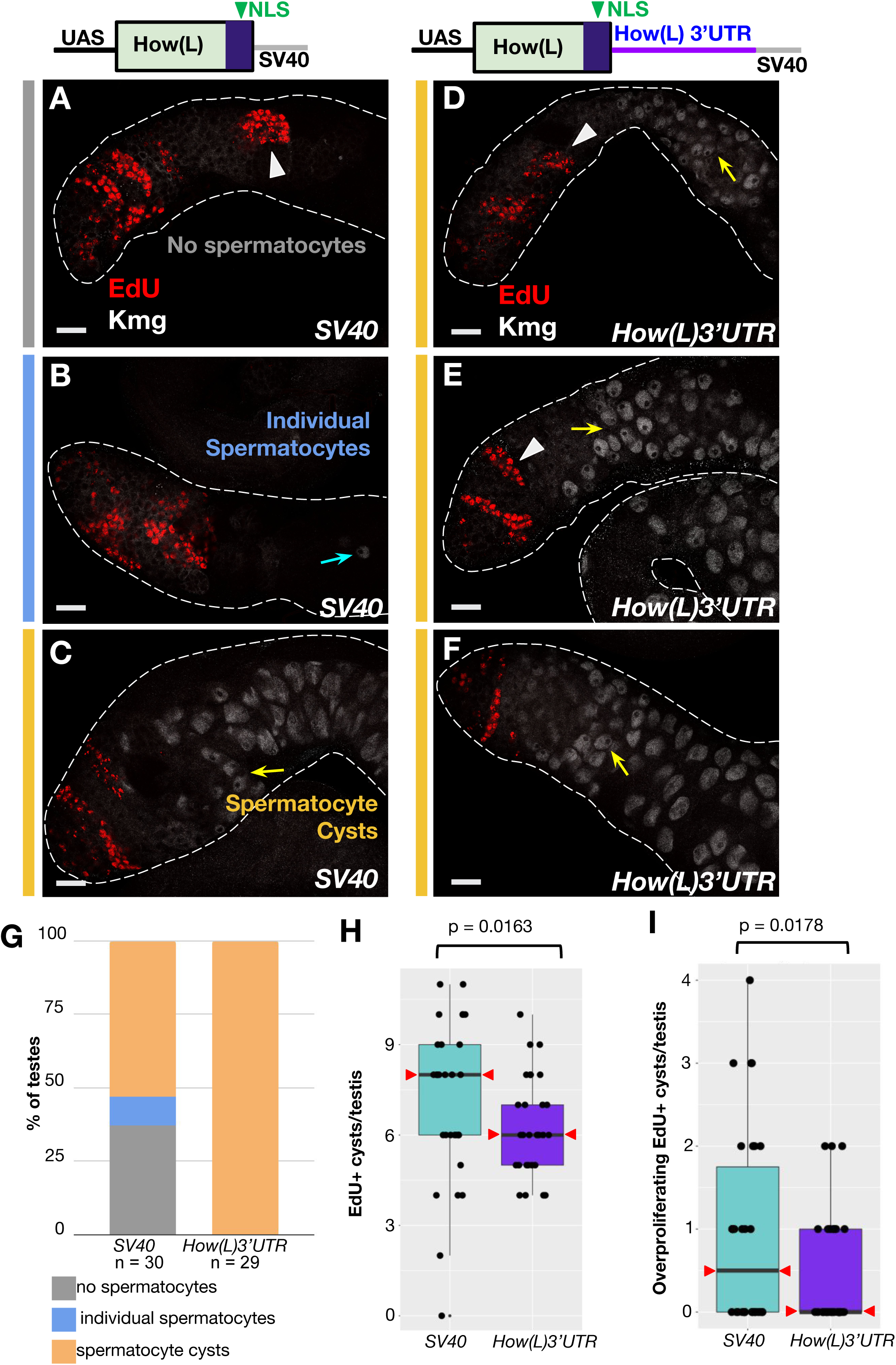
Adding the How(L) 3’UTR reduced the effect of forced expression. (A-F) Top: cartoons depicting design of the overexpression constructs. (Left) *UAS-How(L)HA* with the SV40 terminator, (right) *UAS-How(L)HA* with the 2424nt 3’UTR from How(L) (Figure S2A) added before the SV40 terminator. Immunofluorescence images of testes stained for (white) Kmg to mark spermatocyte nuclei and (red) EdU to label cysts in S phase in the phenotypic classes indicated in G. (A-C) *bamGal4* > *UAS-How(L)HA-SV40*. (D-F) *bamGal4* > *UAS-How(L)HA-3’UTR-SV40*. Flies were grown at 18°C for their entire life. Arrowheads: EdU positive cysts. Blue arrow: single Kmg positive spermatocyte. Yellow arrows: spermatocytes in cysts. Scale Bars in A-F: 25 μm. (G) Graph of the percentage of testes from each genotype with either (gray) no spermatocytes detected, (blue) single spermatocytes or (orange) cysts of multiple spermatocytes, scored based on the spermatocyte marker Kmg. *SV40* n = 30 testes. *How(L)3’UTR* n = 29 testes. Percentages were compared using a two-sample t-test of proportions. Percentage of testes containing spermatocyte cysts (p = 2.5 x 10-5) and percentage of testes containing any spermatocytes (p = 3.0 x 10-4) differed significantly between the two genotypes. (H) Distribution of the number of EdU positive cysts in *bamGal4* > *UAS-How(L)-SV40* testes and *bamGal4* > *UAS-How(L)-3’UTR-SV40* testes. Statistical tests performed via 1-way between-groups ANOVA. (I) Number of overproliferating (>16 nuclei) EdU positive cysts from testes overexpressing *How(L)-SV40* (n = 30 testes) compared to *How(L)-3’UTR-SV40* (n = 29 testes). P-value calculated as for (H). The median in *How(L)-3’UTR-SV40* was zero overproliferating cysts/testis. Medians marked by red arrowheads in all conditions.

Flies bearing either the newly constructed *UAS-How(L)HA-How(L)3’UTR-SV40* transgene (hereafter termed *UAS-How(L)-3’UTR*) or a parallel *UAS-How(L)-SV40* transgene lacking the *How(L) 3’UTR* (hereafter termed *UAS-How(L)-SV40*) inserted at the same genomic site were then crossed to flies bearing *bamGal4* to drive expression starting in mid-to-late spermatogonia and the progeny grown continuously at 18°C. Under these conditions, forced expression of *UAS-How(L)-SV40* had a range of phenotypes, visualized and scored by EdU labeling and anti-Kmg staining. Some testes (37%) had no spermatocytes detected at all (Figure 4A), while the remaining 63% of testes contained at least some individual Kmg-positive spermatocytes (Figure 4B), with 53% of the testes scored containing entire cysts of Kmg-positive spermatocytes (n = 30 testes) (Figure 4C,G). However, in flies in which the *UAS-How(L)-3’UTR* construct was forcibly expressed under control of *bamGal4* at 18°C, 100% of testes had at least some spermatocyte cysts (n = 29 testes) (Figure 4D-F,G). Briefly incubating testes in EdU to label nuclei in S phase revealed that flies overexpressing How(L)-HA with and without the How(L)3’UTR both contained spermatogonial cysts that had undergone additional rounds of proliferation beyond the normal 4, visualized as EdU positive cysts with many more than 16 labeled nuclei (Figure 4A-F,H). However, *UAS-How(L)-3’UTR* testes had overall less spermatogonial overproliferation, based on fewer cysts undergoing S phase per testis (Figure 4H) and fewer cysts with more than 16 EdU labeled nuclei per cyst (Figure 4I), compared to testes from flies overexpressing *UAS-How(L)-SV40* (without the How(L) 3’UTR) grown in parallel. The milder effect of forced overexpression of *How(L)-3’UTR* compared to *How(L)-SV40* under control of *bamGal4* raised the possibility that Bam may downregulate How expression via the How(L) 3’UTR.

### Bam may downregulate *how* by recruiting the CCR4-NOT complex

The reduction of *how* transcript levels in transit amplifying spermatogonia soon after the appearance of Bam protein in the cytoplasm (Figures 1G; S2C-E) suggested that Bam may downregulate *how* at least in part due to effects on the *how* RNA. Protein structure studies by Sgromo *et al.* (2018) showed that a 23 amino acid sequence near the N-terminus of the Bam protein binds in a groove of Caf40, a subunit of the CCR4-NOT complex (10). Further, Sgromo *et al.* showed that when Bam or an N-terminal fragment of Bam containing the Caf40-binding motif (CBM) was tethered to a luciferase reporter RNA, it was able to recruit CCR4-NOT for transcript degradation, decreasing levels of luciferase RNA and protein (10). These findings raise the possibility that Bam and Bgcn may downregulate *how* by recruiting Caf40. Consistent with this, knockdown of *Caf40* in early spermatogonia by RNAi under control of *nosGal4* resulted in massive overproliferation of small germ cells, similar to the phenotype observed in *bam* mutant males (Figures 5C; S5B-E). Digesting the testis sheath to spill out intact cysts confirmed that the overproliferating small cells were organized in cysts (Figure 5D), as in *bam* mutants. Brief incubation in EdU confirmed that large clusters of germ cells were undergoing DNA synthesis in synchrony in testes in which expression of *Caf40* had been knocked down under control of *nosGal4*, while immunofluorescence staining confirmed failure to turn on expression of the spermatocyte marker Kmg (Figure 5F), as in *bam* mutant males. Strikingly, immunofluorescence staining with anti-Bam revealed that the hundreds of small germ cells that overproliferated when *Caf40* was knocked down by RNAi under control of *nosGal4* expressed Bam protein (Figure 5H), indicating that they were germ cells and that the failure to differentiate was not due to failure to express Bam.

**Figure 5.**
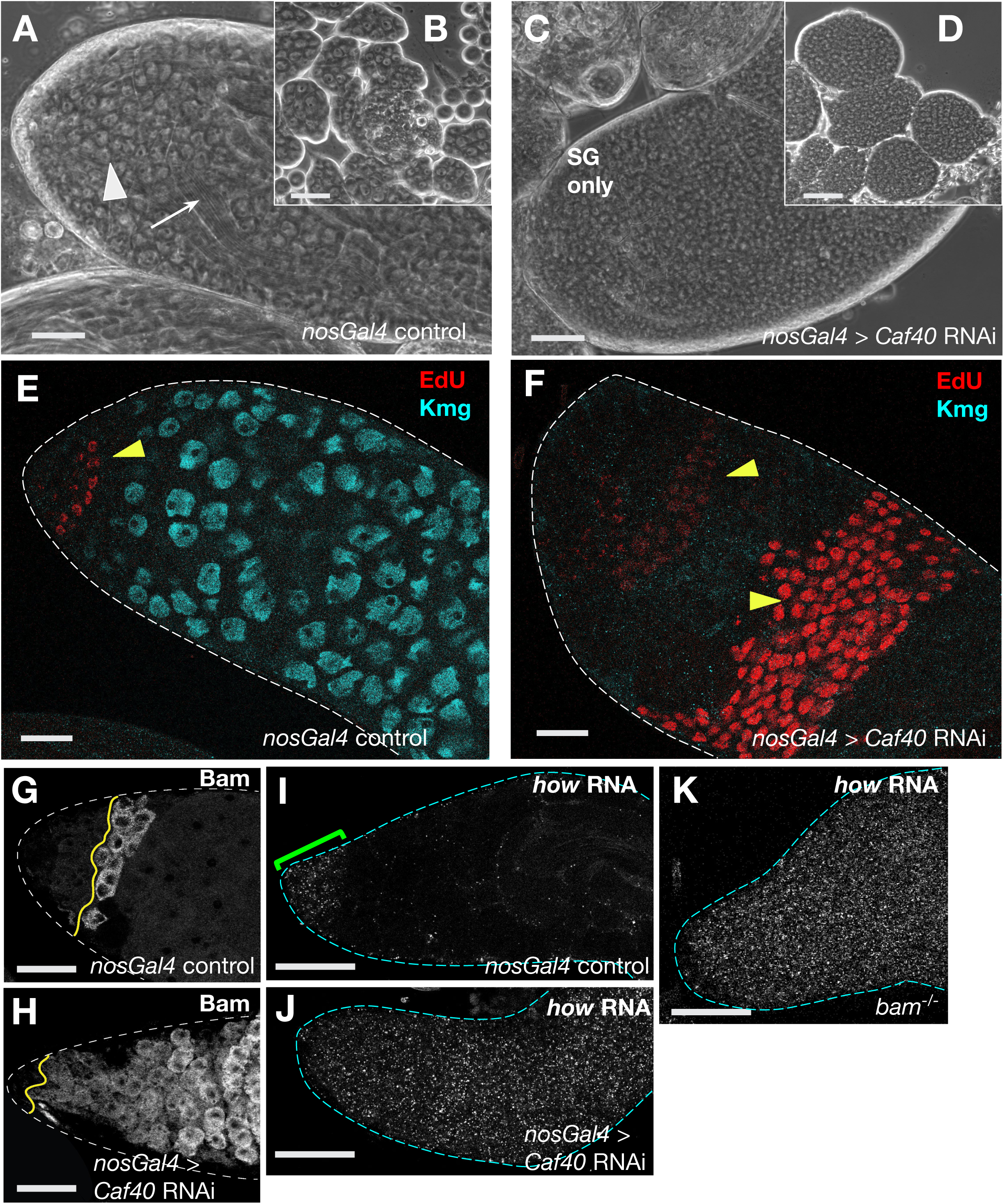
The CCR4-NOT component Caf40 is required for *how* repression. (A, C) Phase contrast images of testis apical tips from males carrying (A) *nosGal4* driver only control and (C) *nosGal4* > *Caf40 RNAi* (TRiP HMS05850 Bloomington stock #67987). White arrowhead: spermatocytes. White arrow: spermatid bundles. SG = spermatogonia. (B, D) Spilled out cysts from (B) *nosGal4* control testes including many spermatocyte cysts as well as more uncommon cysts of smaller spermatogonia. Spilled out cysts from (D) *nosGal4* > *Caf40* RNAi males contained many small cells and no spermatocytes. (E, F) Immunofluorescence images of testes apical tips from (E) *nosGal4* driver only control and (F) *nosGal4* > *Caf40* RNAi labeled with (red) EdU to mark nuclei in S phase and (cyan) anti-Kmg to mark spermatocyte nuclei. Yellow arrowheads: cysts of EdU positive spermatogonia. (G, H) Immunofluorescence images of testis apical tips stained with anti-Bam from (G) *nosGal4* and (H) *nosGal4* > *Caf40* RNAi flies. Yellow line: boundary between early spermatogonia and Bam positive spermatogonia. (I-K) Apical tips of testes stained for *how* RNA by HCR with probes complementary to the How protein coding sequence. (I) *nosGal4* control. Green bracket: region with *how* transcripts in spermatogonia. (J) *nosGal4* > *Caf40* RNAi. (K) *bam^1/^*^Δ*86*^. Scale bars: 50 μm A-D; 25 μm E-K. All control flies were raised under the same temperature regimen used for the RNAi (3 days at 25°C, then cultured at 29°C).

Analysis by fluorescence HCR *in situ* hybridization (FISH) confirmed that *how* transcripts remained in the germ cells that overproliferated after function of the CCR4-NOT component *Caf40* was knocked down by RNAi. In testes from control males carrying the *nosGal4* driver but not the *UAS-Caf40 RNAi* construct (Figure 5I) raised under RNAi temperature shift conditions, *how* mRNA was detected in early germ cells near the testis apical tip but was not detected in the spermatocyte region further down the testes (as in Figure 1).

However, despite the expression of Bam protein, *how* transcripts remained high in the overproliferating germ cells that accumulated in testes in which *Caf40* had been knocked down in early germ cells under control of *nosGal4*, as observed in *bam^-/^*^-^ males (Figure 5J,K). The persistence of *how* transcripts despite the presence of Bam protein suggested that Caf40 does not act upstream of Bam but instead might be part of the mechanism through which Bam protein downregulates *how* RNA.

## Discussion

### A key function of Bam in spermatogonia is repression of *how*

Our results indicate that the major role of Bam and Bgcn in the switch from mitotic proliferation to onset of the meiotic program in the male germ line is to down regulate expression of Held Out Wings, the *Drosophila* homolog of mammalian Quaking (Figure 6A). Most telling, male germ cells lacking *bam* function can differentiate if expression of *how* is knocked down in mid-to-late stage transit amplifying spermatogonia by RNAi. In addition, forced expression of a nuclear targeted isoform of How, How(L), was sufficient to cause spermatogonia to overproliferate and largely fail to become spermatocytes. Strikingly, spermatogonia subjected to forced expression of How(L) expressed abundant Bam protein, indicating that the failure to differentiate was not due to How(L) repressing expression of Bam.

**Figure 6.**
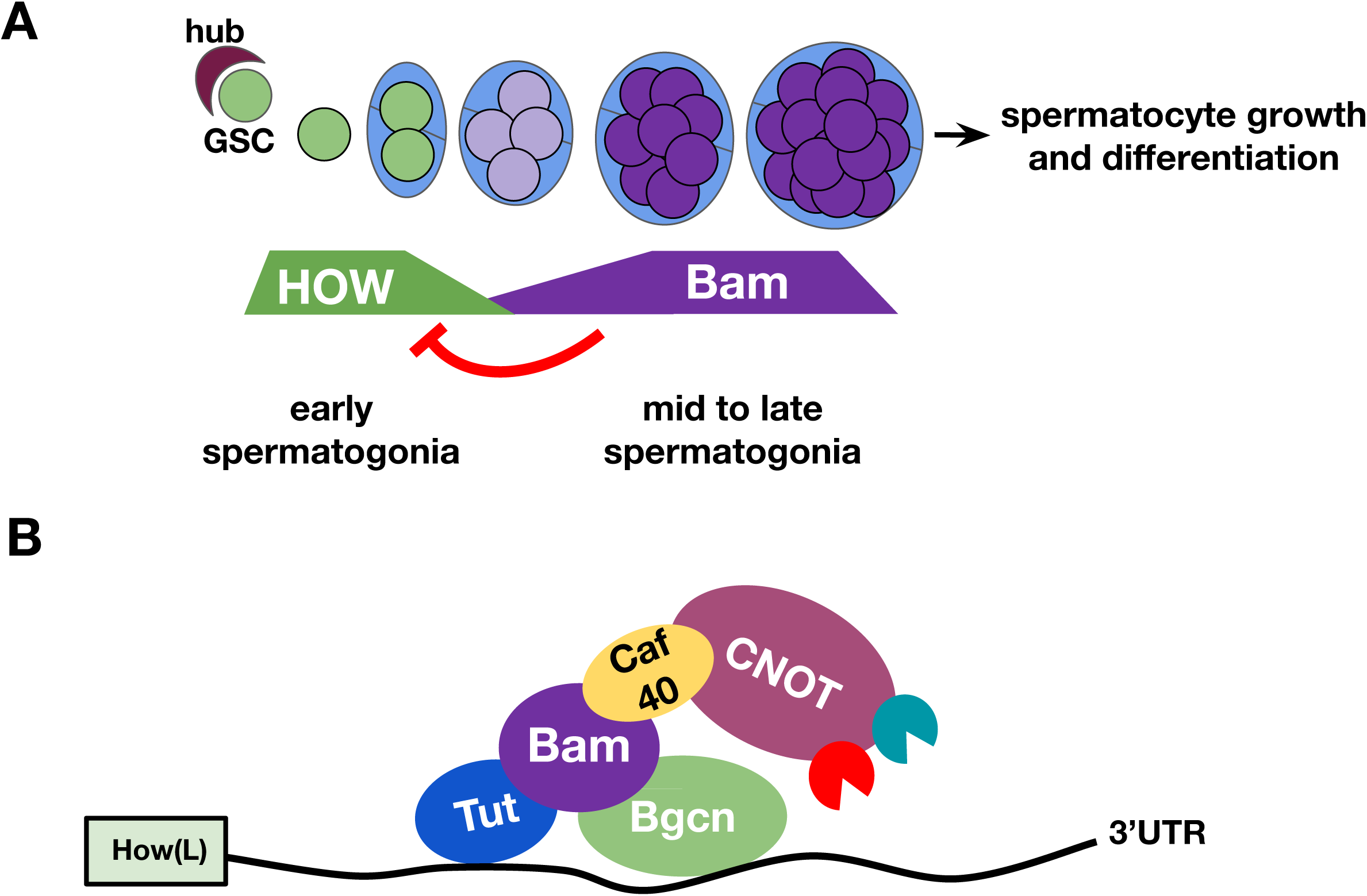
Model for mechanism of Bam action on *how* RNA. (A) Cartoon of *How* (green) and *Bam* (purple) protein expression in the germ line stem cell and spermatogonia. Levels of *how* decrease as Bam expression appears at the four cell cyst stage. Bam levels increase in mid-to-late spermatogonia until sufficient to repress expression of *How*, allowing the subsequent switch from proliferation to differentiation. (B) Model for regulation by Bam, Bgcn and Tut: Bam protein binds the CCR4-NOT subunit Caf40, acting as an adaptor between the CCR4-NOT complex and its target transcripts.

Our finding that knockdown of *Caf40* by RNAi driven by *nosGal4* led to overproliferation of spermatogonial cysts and persistence of *how* RNA and protein, even though Bam protein was present, suggested that *Caf40* is required for Bam to downregulate *how*. Sgromo *et al.* showed that Bam protein has a 23-amino acid N-terminal domain that binds in a groove of *Caf40* (homolog of human NOT9), a subunit of the CCR4-NOT complex (10). Through this Caf40-binding motif (CBM) domain, Bam tethered to a reporter RNA was able to recruit the CCR4-NOT complex to degrade the reporter RNA (10). Bam protein appears to participate in a ternary complex, bridging between the RNA binding protein Tut bound to sequences in the Bam N-terminal third and the RNA binding protein Bgcn bound to sequences in the Bam C-terminal third (8). Consistent with models suggested by others, we propose that Bam, recruited to target RNAs by its structural partners Bgcn and Tut, may act as an adaptor to recruit the CCR4-NOT deadenylation complex to destabilize the bound RNA and/or lead to its translational repression (8, 10, 16) (Figure 6B). In support of the model that Bam targets *how* RNA for degradation and/or translational repression by recruiting CCR4-NOT, knockdown of several CCR4-NOT subunits by RNAi under control of the *bamGal4* driver showed a mild overproliferation phenotype in spermatogonial cysts (16).

Bam and Bgcn have been shown to downregulate expression of E-Cadherin via sequences in the E-Cadherin 3’UTR (17). Assays conducted in S2 cells carrying reporter constructs with the Firefly *luciferase* coding sequence attached to the 3’UTR from E-Cadherin showed that expression of luciferase was down regulated compared to a renilla control with a heterologous 3’UTR if Bam and Bgcn were co-expressed in the cells. The inhibition of luciferase expression was not observed when either Bam or Bgcn was expressed without its partner, but was detected if Bam was tethered to the E-Cadherin 3’UTR. Bgcn, Bam and Tut have also been shown to repress expression of Mei-P26 via sequences in the *mei-P26* 3’UTR (8, 11).

Bam serves as a differentiation factor in both the male and female germ lines (6, 7). The regulatory logic is similar: in both cases, action of *bam* downregulates a factor required for maintenance of the precursor state. In *Drosophila* females, the germ line stem cell state is maintained by *nanos* and *pumilio*, which are thought to repress translation of transcripts that drive differentiation (7, 18–20). Loss of function of *nanos* or *pumilio* in females resulted in loss of germ line stem cells to differentiation (19). In female early germ cells, expression of *nanos* protein was abruptly downregulated when Bam protein became expressed in late cystoblasts. Nanos protein expression persisted in the oogonia that overproliferate in *bam* mutant ovaries, and the down regulation of *nanos* in response to Bam depended on sequences in the *nanos* 3’UTR (7, 21). Similarly, in males, How protein is downregulated when Bam becomes expressed in mid stage transit amplifying spermatogonia, How protein persisted in spermatogonia mutant for *bam*, and addition of the How(L) 3’UTR somewhat reduced the effects of forced expression of How(L).

The timing of Bam expression and action is different in male than in female germ cells. In females, Bam protein becomes upregulated enough to be detected by immunofluorescence staining in the late cystoblast, the immediate daughter of a stem cell division, which will found a clone of female germ cells that will eventually generate one oocyte and fifteen nurse cells. In males, Bam protein is upregulated later, during the spermatogonial transit amplifying divisions, so that it was initially detected by immunofluorescence staining in four cell cysts, appeared to increase in level by the 8 cell stage, and remained present through pre-meiotic S phase before being abruptly degraded (4). In both sexes, expression of How protein in germ cells appears reciprocal to expression of Bam. In females, How protein was present in the nucleus of female germ line stem cells but was abruptly down regulated by the two cell stage (22). Male germ line stem cells also showed nuclear How, which persisted in gonialblasts and two cell transit amplifying stages but dropped precipitously once Bam protein was expressed and began to accumulate (12, this study).

In males, knockdown of *how* in germ line stem cells and early transit amplifying cells under control of *nosGal4* led to loss of germ line stem cells, as did induction of germ line clones homozygous for a strong loss of function *how* allele (12). Germ cells homozygous mutant for *how* appeared stalled in G2 at the two cell cyst stage, likely due to defects in expression of *Cyclin B*, and eventually died (12). The effect of loss of How function in female germ line stem cells was different: induction of germ line clones mutant for *how* led to gradual loss of female germ line stem cells, possibly to differentiation, but not to cell cycle arrest and apoptosis (22). These results suggest that although the regulatory relationship between How and Bam may be similar, the roles of How (and Bam) in early germ cells in the two sexes likely differs.

Monk *et al*., 2010 (12) observed that using *nosGal4* to drive overexpression of *How(L)-SV40* in early germ cells (approximately the time when How is normally expressed) caused shortening of cell cycle time in TA spermatogonia, likely due to increased expression of Cyclin B, and frequent 32 cell spermatocyte cysts, indicating switching from spermatogonia to spermatocytes after 5 rather than the usual 4 rounds of transit amplifying mitotic divisions. Monk *et al.*, 2010 proposed that How(L) represses expression of Bam protein in early spermatogonia (12). In contrast, we found that using *bamGal4* to drive forced expression of *How(L)-SV40* later, a time when it is normally downregulated, resulted in massive overproliferation of spermatogonia. Strikingly, Bam protein was expressed in the spermatogonia overproliferating under these conditions, arguing that How(L) does not repress expression of Bam in later stage spermatogonia.

Our data suggest that Bam down regulates expression of How in mid-to-late transit amplifying spermatogonia, possibly through the How(L) 3’UTR, which was not included in the *How(L)-SV40* construct. Such a mutual repression relationship may provide a mechanism to convert a gradual rise in Bam expression (perhaps driven at the level of transcription) into a sharp switch in cell state. In early transit amplifying male germ cells, abundant How(L) protein may repress premature accumulation of Bam protein and maintain a stem cell competent, early TA state. As expression *of bam* transcript and protein rise, however, Bam may reach levels sufficient to bind via a Tut-Bam-Bgcn complex to all or most of the How(L) mRNA molecules, targeting them for translational repression and/or degradation and so throwing the switch to onset of differentiation. Alternatively, the 32 spermatocyte cell cysts detected by Monk *et al.,* (2010) could be due to the observed shortening of the cell cycle in spermatogonia after forced expression of How(L) under control of *nosGal4* (12) rather than to How(L) repression of *bam* in early spermatogonia. As shown by Insco *et al.* (2009), shortening the cell cycle can allow more cysts to complete 5 rounds of cell division before Bam protein reaches the critical level required for the switch to spermatocyte state (4).

The fact that testes from *bamGal4>UAS-How(L)-3’UTR* males still showed some aberrant overproliferation of spermatogonia may be because getting the proportions of *how* RNA and Bam protein just right so that all the *how* RNA can be bound and repressed/degraded is not easy. The amount of How(L)-3’UTR mRNA produced in the *bamGal4>UAS-How(L)-3’UTR* males may titrate out the endogenous Bam protein and so overwhelm the capacity of Bam to repress either the *UAS-How(L)* or endogenous *how* RNA.

A central lesson from genetic analysis of the *Drosophila* male and female germ line adult stem cell lineages is that the switch from precursor cell state to onset of differentiation is controlled by a cascade of RNA-binding proteins. In each case, Bam protein with its RNA-binding partner Bgcn acts to trigger differentiation by downregulation of another RNA-binding protein. In females, the key target is the translational inhibitor *nanos* (7). Here we show that in males the key target is the RNA-binding protein How. It will be interesting in future work to test whether mRNAs bound and regulated by How(L) in early spermatogonia maintain the germ cells in a proliferative state or block their differentiation. Our results suggest that it will also be important to investigate whether similar regulation by RNA-binding proteins controls the switch from proliferation to differentiation in adult stem cell lineages in other organisms.

## Methods

### Fly Strains and husbandry

Flies were raised on molasses food. Overexpression and knockdown crosses were raised at 25°C for 3 days, then shifted to 29°C after discarding adults, unless otherwise noted. RNAi lines were obtained from The Vienna Drosophila Resource Center (VDRC): UAS *how* RNAi (100775), UAS *caf40* RNAi (101462) and the Bloomington Drosophila Stock Center: UAS *how* RNAi (Bloomington stock #55665, containing TRiP line HMC03820) and UAS *caf40* RNAi (67987, containing TRiP line HMS05850). Results using the How RNAi line VDRC100775 are shown throughout except for Figures 2I, S3, and S6, which also show results from knocking down How using the TRiP RNA construct HMC03820 carried in Bloomington stock #55665.

For Figure 4, flies were raised at 18°C throughout. For the heat shock time course, flies were raised at 25°C until there were mid- and late-stage pupa. Then bottles were placed in a 37°C water bath for 30 minutes to induce heat shock and returned to 25°C, as spelled out in Kim *et al*., (2017) (13). For the RNA-Seq analysis shown in Figure S5, the *bam* mutant testes were from *CyO/+; bam^1^*/*bam*^Δ86^ males that were grown at 22°C until the late pupal stage, then the bottles were incubated in a 37°C water bath for 30min, then cultured at 25°C for 48hrs, at which time the testes plus seminal vesicle were obtained by dissection and prepped for RNA extraction. In all other cases, flies were raised at 25°C. *UAS-How(Long)-3xHA* and *UAS-How(Short)-3xHA* fly stocks with an SV40 heterologous 3’UTR were a gift from T. Volk (23). Two Gal4 driver lines were used: *nanosGal4*-*VP16* for germ line stem cells and early spermatogonia and *bamGal4* for the mid-to late spermatogonia (24, 25). Gal4 driver strains for knockdowns also contained *UAS-Dicer2*. The *bam* alleles were *bamP*-*bam-GFP* (25), *bam^1^*(5), and *bam*^Δ86^ (26). Fertility tests were performed with males from crosses that had been shifted to 29°C for Gal4 driver expression. One male was placed in a vial with three virgin females and kept at 25°C. Adults were removed after six days and vials were scored for the presence of pupa and adult offspring after at least 10 days.

### Phase Contrast Microscopy

For testis squashes, testes from 0-1 day old males were dissected in PBS. Whole testes or cysts were flattened into a monolayer under a coverslip by wicking away PBS and observed by phase contrast microscopy using a Zeiss Axioskop. Images were taken with a Spot Imaging camera and software. To count the number of spermatocytes per cyst, dissected testes were treated with 0.5 mg/mL collagenase (Sigma C7657) + 0.5 mg/mL dispase (Worthington, LSO2109) in PBS on a slide for 1.5-3 minutes (larger testes burst open sooner) (as described in Lu *et al*., 2020) (27). The reaction mix was replaced with PBS and cysts were gently flattened under a coverslip by wicking away liquid before counting and imaging.

### Transgenics

Plasmids containing *how* coding sequences with HA tags expressed under control of UAS were a gift from Talila Volk (23). The *how*(L) coding sequence (FBgn0264491, isoform A on FlyBase) was amplified and inserted into a pUASTattB vector at NotI and XhoI restriction sites (15). To add the *how*(L) 3’UTR ( FBtr0084177) was amplified in two parts from *bam* testis cDNA (to exclude the intron) and inserted upstream of the SV40 polyadenylation site, using XhoI and XbaI restriction sites. Differences in the *how* coding sequence from published fly genome (Dm6) were corrected to match *bam^1^*/*bam*^Δ86^ mutant flies, using a Q5 Site-Directed Mutagenesis Kit (New England Biosciences E0554S). Plasmids were injected into PhiC31 integrase transgenic flies and the constructs were integrated at attP40 on chromosome 2L, then selected for transformants by BestGene Inc. (Chino Hills, CA) (28).

### Immunofluorescence

For whole mount preparations, whole testes were dissected in PBS, fixed in 4% paraformaldehyde (PFA) for 20 minutes and washed twice in PBS. The tissue was permeabilized in a solution of 1X PBS, 0.6% sodium deoxycholate, and 0.6% Triton for 1 hour at room temperature, then washed twice before blocking overnight in PBST(Triton)-3% BSA (bovine serum albumin) at 4°C. Primary antibodies were added at the following concentrations and testes were incubated rotating for two days: rabbit anti-HOW (1:50), mouse anti-Bam from DSHB (1:10), rabbit anti-Kmg (1:200) (Kim et al., 2017, 13), mouse anti-HA (1:200), and goat anti-Vasa (1:100, Santa Cruz Biotechnology). After two PBS washes at room temperature, donkey secondary antibodies (Alexa Fluor, ThermoFischer) were added at 1:500 and the samples were incubated for 2 hours at room temperature. Testes were then mounted on a slide in DAPI mount (Vectashield). To decrease background/non-specific binding, the rabbit anti-HOW antibody (gifted by T. Volk) was preabsorbed with ∼10 new wild-type testes overnight 3 times (fixed as for whole mount preparations.

For squashed preparations (samples stained with anti-HOW and anti-Bam), testes were placed on a SuperFrost Plus slide in a square drawn with a hydrophobic marker. Then the tissues were flattened live under a coverslip before being flash frozen in liquid nitrogen. After removing the coverslip, slides were incubated in cold 95% ethanol for 10 minutes before being fixed in 4% PFA for 7 minutes. Testes were washed in PBST (Triton-X) before being transferred to a wet chamber for blocking overnight in BSA at 4°C, after which antibody staining continued as for whole mounts.

### EdU Labeling

Cells in S phase were labeled with the Click-iT EdU Cell Proliferation Kit for Imaging - Alexa Fluor 555 dye (Invitrogen - C10338). Testes were dissected in Schneider’s (S2) medium and transferred into a drop of the same medium on a slide. The medium was then removed and replaced with 100 μM 5-ethynyl-2′-deoxyuridine (EdU) in S2 cell medium and incubated for 5 minutes at room temperature. Testes were then washed twice in S2 medium by removing the liquid with a pipette and replacing it with S2 medium. After the washes, testes were transferred with tweezers from the drop of S2 medium into a 1.7 mL Eppendorf tube with ∼200 uL 1xPBS, then immediately the PBS was drawn off and replaced with 200 μL 4% paraformaldehyde in PBS for 20 minutes (rotating at room temperature) to fix, followed by two washes in PBST. The testes sheaths were then permeabilized in PBS with 0.6% Triton and 0.6% sodium deoxycholate for one hour. The permeabilization mix was removed and the testes were rinsed once with PBST before adding the EdU detection reaction mix per the manufacturer’s instructions. Testes were then incubated in the dark with the reaction mix for 30 minutes, the reaction mix was removed, and the testes washed twice in 500 μL of PBST at room temperature. Blocking and antibody protocols continued in the same way as for squashes and whole mounts.

### Hybridization chain reaction *in situs*

Probes to label the *how* coding sequence and the *how(L) 3’UTR* were designed following the method of Bedbrook et al., 2023 (29), which generated 44 probes (22 pairs) then ordered from Integrated DNA Technologies at 50 pmol/oligo. All other reagents were from Molecular Instruments (Los Angeles, CA), including H1 and H2 hairpins conjugated with 488. Testes were dissected in 1xPBS, fixed in 4% paraformaldehyde for 20 minutes, then permeabilized in 0.6% sodium deoxycholate for 30 minutes. Samples were then washed twice in PBS at RT.

Hybridization chain reaction *in situ* hybridization followed the “sample in solution” HCR™ RNA-FISH protocol from Molecular Instruments with the following specifications: Probe solution was made with 1 μM of probes. Hybridization buffer and probe wash buffer were reduced to 200 μL from 500 and pre-amplification was done in 250 μL of amplification buffer. After completing the HCR protocol, DAPI mounting media was added, testes were mounted on a slide and imaged by confocal microscopy.

### Imaging and quantification

Immunofluorescence stained or HCR FISH testes were imaged on a Leica SP8 Confocal microscope. Image brightness was adjusted in FIJI. To quantify overproliferation, testes were scored for the overall number of EdU positive cysts per testis and the number of EdU positive cysts with >16 cells each. Testes were also stained for the spermatocyte marker Kmg and scored for the presence of individual Kmg positive spermatocytes or spermatocyte cysts. For the quantification in Figure 4, EdU and Kmg scoring was performed with the scorer blind to the genotypes.

### Analysis of transcripts expressed in testes

Microarray experiments were performed as described in Kim *et al.,* 2017 (13). To control for possible persistent changes in gene expression resulting from the heat shock treatment, control *bam^1^*/*bam*^Δ86^ flies lacking the *hsBam* transgene were subjected to 30min of heat shock then cultured at 25°C for 8, 16, 24, or 32hr in parallel with the experimental *hsBam; bam^1^*/*bam*^Δ86^ flies and the genes expressed in testes from the two genotypes were assessed by microarray in parallel. Briefly, RNA was extracted from about 30 pairs of dissected testes with seminal vesicles but without accessory glands using Trizol (Invitrogen). Reverse transcription was performed with oligo(dT)24 primer with a T7 promoter using ∼200 ng of total RNA per Affymetrix protocol. The second strand was synthesized from the cDNA, and cRNA was produced by *in vitro* transcription. Fragmented cRNA was hybridized to the *Drosophila* genome 2.0 arrays (Affymetrix, Cat# 900532). All microarray experiments were performed by the Stanford Protein and Nucleic Acid (PAN) facility. *how* transcript isoforms were distinguished by probes binding at the 3’ end of transcripts, identified by Affymetrix Probeset ID (1637943_at). For analysis, all the raw CEL files were background adjusted and quantile normalized together by using R/BioConductor (v3.0.2) package GCRMA (30). Gene annotation was based on the Affymetrix file: “Drosophila_2.na32.annot.csv”.

For analysis of transcript expression by RNA-Sequencing, 10∼20 μg of total RNA was extracted from ∼100 pairs of testes plus seminal vesicle from *hsBam; bam^1^*/*bam*^Δ86^ flies for each time point using Trizol (Invitrogen) followed by RNeasy cleanup (Qiagen) according to kit instructions. PolyA-tailed RNA was purified using the Oligotex mRNA kit (Qiagen, Cat#70022). Purified PolyA RNA was fragmented in the presence of random hexamer primers in first strand synthesis buffer (Invitrogen, Cat# 18080093) at 85°C for 8 minutes. Fragmented RNAs were reverse transcribed using Superscript III reverse transcriptase (Invitrogen, Cat# 18080093) in the presence of RNaseOUT (Invitrogen Cat# 10777019) at 50°C for one hour.

From this step, we followed the directions in the NEBNext mRNA Library Prep Master Mix Set for Illumina (E6110s) to make libraries. DNA was purified using a QIAquick PCR purification kit (Qiagen Cat# 28104) after second strand synthesis, end repair, dA tailing, and adapter ligation. After adapter ligation, 300∼500 bp fragments were size-selected by gel extraction (1.5% low-melt NUSIEVE gel in TBE). Pooled libraries were sequenced with Illumina HiSeq: 100bp, each paired-end with single indices.

For analysis of time course RNA-Seq data, raw fastq reads were trimmed using trim galore (version 0.4.3) to remove low-quality (Phred score 20) and adapter-containing (stringency 1) reads (31). Trimmed reads were mapped to the *Drosophila melanogaster* genome (BDGP6.46) with default parameters using STAR (version 2.5.3) (32). Mapped reads with quality scores smaller than 10 (-q 10) or not properly paired reads (-f 2) were discarded using SAMtools (version 1.4.1) (33). Counts per gene were obtained using the featureCounts function of the subread package (version 1.5.0) (34). Expression levels at different time points were TMM-normalized (Trimmed Mean of M-values), assuming the majority of housekeeping genes have the same expression levels in different time points using the R (version 4.1.0) package edgeR (version 3.34.0) (35). Axes for scatter plots were log2 transformed normalized CPM (Counts Per Million) + 1. Scatter plots were generated by Matlab (R2021a). Analyses scripts are available in: https://github.com/jongminkmg/HeldOutWings2024.

To ensure knockdown of *caf40,* we analyzed gene expression levels using RNA-seq for testes plus seminal vesicle from *bam* mutant (48hrs after heat shock) and *nosGal4* driven *caf40* knockdown flies (Figure S5). Testes were dissected from 0–2 day old males in 1xPBS, as batches of 50 flies, in a cyclops dissecting dish for <30min at room temperature. Each batch was transferred to a 1.7 ml Eppendorf tube containing 1ml of 1xPBS, the PBS was then immediately removed, and the testes were snap-frozen in liquid nitrogen and stored at −80°C.

RNA was extracted from 150 pairs of testes using the RNeasy Plus Mini Kit from QIAGEN (74104). Frozen tissue was dissociated using a 1 ml syringe with a 27-gauge needle aspirating up and down ∼10 times in 300µl of lysis buffer (from RNeasy kit) supplemented with 1:100 β-mercaptoethanol. Library preparation was carried out using the NEBNext® Ultra™ II Directional RNA Library Prep Kit for Illumina (#E7760S), with the NEBNext® Poly(A) mRNA Magnetic Isolation Module (#E7490S). Sequencing was performed by Novogene on a NOVAseq, PE 150 Illumina platform. Adapters and low-quality bases were trimmed with trimGalore and then aligned to the *Drosophila* dm6 genome using STAR. Differential expression analysis was performed using DEseq2.

## Acknowledgements

We thank members of the Fuller Lab for their insights on the project and helpful suggestions and T. Templin for help with statistical analysis. DH thanks Anne Villeneuve, Lauren Goins, and Roel Nusse for their guidance on experiments, interpreting data, and data visualization. Fly food and solutions were made by The Developmental Biology Department Fly Food Facility and Media Facility. Talila Volk generously provided UAS-HOW fly stocks and How antibodies. The Stanford Protein and Nucleic Acid (PAN) facility assisted with the microarray experiments. Kitty Lee and Jon Mulholland of Stanford’s Cell Sciences Imaging Facility (CSIF) provided invaluable training and assistance on the Leica SP8 Confocal Microscope.

This work was funded by NIGMS MIRA #1R35GM136433, the Katherine Dexter McCormick and Stanley McCormick Memorial Professorship, and the Reed-Hodgson Professorship in Human Biology to MTF, the Stanford University Genetics and Developmental Biology Training Grant 5T32GM007790 (DH), and The Italian American Cancer Research Fellowship (LG). RNA-sequencing from the heat shock time course was partially funded by 1U54HD068158 (PI: Renee Reijo Pera). The project described was also supported, in part, by Award Number 1S10OD010580-01A1 from the National Center for Research Resources (NCRR). Its contents are solely the responsibility of the authors and do not necessarily represent the official views of the NCRR or the National Institutes of Health.

## SOM Figure Legends

**Figure S1. *how* is among the earliest transcripts to decrease after Bam is turned on.**

(A, B) Scatterplots of transcript levels from microarray analysis comparing no heat shock and 8 hours PHS in (A) *bam^1/^*^Δ*86*^; *hs-Bam* or (B) *bam^1/^*^Δ*86*^ males lacking the *hs-Bam* construct, but subjected to the same heat shock regimen in parallel, to control for the effects of heat shock on gene expression. Genes colored blue (downregulated) or red (upregulated) were 1) not changed >2 fold in the microarray analysis of testes from control *bam^1/^*^Δ*86*^ males lacking the *hs-Bam* construct but heat shocked then incubated for the indicated time and 2) also detected by independent RNA-seq as up or down >2 fold in *bam^1/^*^Δ*86*^; *hs-Bam* males by RNA-sequencing at the time points indicated compared to testes from flies of the same genotype not subjected to heat shock. (C) Level of *how(L)* transcripts detected by microarray throughout the time course, showing decrease by 8h PHS in flies carrying the *hs-Bam* construct, but not in testes from control *bam^1/^*^Δ*86*^ flies lacking the hs-Bam construct but subjected to the 30-minute pulse of incubation at high temperature, then shifted back to 25°C as for the experimental genotype. (D, E) Microarray data from later time points after heat shock, showing the transcripts that increase in expression (red) and decrease in expression (blue) by the cutoff criteria (detected as up or down regulated >2 fold) in both this microarray comparison and in independent analysis by RNA-seq. (D) *bam^1/^*^Δ*86*^; *hs-Bam* testes (red oval in *hs-Bam* 32h: genes expressed specifically in spermatocytes. (E) testes from control *bam^1/^*^Δ*86*^ males that did not have the *hs-Bam* transgene, but were subjected to heat shock then incubated at 25°C for the indicated times. Black oval in (E) 16h PHS marks genes expressed in accessory glands that contaminated this sample.

**Figure S2. How transcripts were detected in spermatogonia but not in spermatocytes**

(A) Diagram of the *how* locus showing mRNA isoforms from FlyBase, designated A-F as in Flybase. The How(L) cDNA construct utilized in Figures 3 and 4 is RA and the How(S) cDNA construct utilized in Figure 3 is RB. Grey: UTRs. Green: protein coding sequence. Lines denote introns. Red arrowheads: nuclear localization signal. Orange arrowhead: additional coding sequence in isoform RF, distinguishing it from RA. Black bars: Sites of RNAi constructs utilized. (B) Apical tip of wild type testis showing distribution of *how* transcripts detected by Hybrid Chain Reaction (HCR FISH) using two different probe sets. (B) Merge. Magenta: Coding sequence (CDS) probe set; Green: 3’UTR probe set. Zoom in marked by red box. Yellow rectangles mark white loci of overlapping probes sets. (B’) How(L) 3’UTR probe set. (B”) Coding sequence (CDS) probe set. Scale Bar: 25 μm. (C-E) High magnification immunofluorescence images of apical tips of *Bam-GFP* testes with How protein coding sequence RNA labeled by HCR (additional examples for Figure 1G). Left: merge with (blue) DAPI, (green) *how* RNA, and (magenta) Bam-GFP. Middle: *how* RNA only. Right: Bam protein only. Yellow dashed outlines: early Bam positive cysts with *how* RNA present. White dashed outlines: later stage Bam positive cysts with fewer h*ow* RNA foci. Cyan dashed outlines: Bam positive cysts with no or very low *how* RNA signal detected. Scale bar: 12.5 μm.

**Figure S3. Knockdown of *how* in mid-to late spermatogonia in *bam* mutant males resulted in production of spermatocytes for two different RNAi lines.**

(A-C) Phase contrast images of testis apical tips from (A) *bamGal4* driver only; (B) *bam^1/^*^Δ*86*^; and (C) *bam^1/^*^Δ*86*^; *bamGal4* > *how* RNAi (TRiP HMC03820 from Bloomington Stock #55665). Arrowheads: spermatocytes. Arrows: spermatid bundles. SG: spermatogonia. Scale bars: 100 μm, insets Scale Bars: 25 μm. (D-F) Immunofluorescence images of apical testis tips stained for EdU to mark S-phase nuclei (red) and Kmg to mark spermatocyte nuclei (blue). (D) Control: *bam*Gal4 driver only. (E) *bam^1/^*^Δ*86*^. (F) *bam^1/^*^Δ*86*^; *bam*Gal4 > *how* RNAi (TRiP HMC03820 from Bloomington Stock #55665). (G-I) Whole testis from (G) *bamGal4* driver only; (H) *bam^1/^*^Δ*86*^; and (I) *bam^1/^*^Δ*86*^; *bamGal4* > *how* RNAi (VDRC100775) for size comparison. Arrowhead: spermatocytes. Arrow: spermatid bundles. Scale bars: 100 μm. (J) Fertility tests of *bamGal4* driver only, *bam^1/^*^Δ*86*^; and *bam^1/^*^Δ*86*^; *bamGal4* > *how* RNAi (VDRC 100775) males. Males from all genotypes were progeny from crosses where the mated parents were allowed to lay eggs for 3 days at 25°C then the parents were removed and the progeny shifted to 29°C. Once the progeny males were collected, they were subjected to fertility tests at RT, ∼ 25°C.

**Figure S4: Knockdown of *how* in mid-to late spermatogonia in *bgcn* mutant males resulted in production of spermatocytes.**

Immunofluorescence images of apical tip of *bgcn^1/63-44^*mutant testis stained with (A) anti-How and (A’) anti-Vasa. Scale bars: 25 μm. (B,C) Phase contrast images of testis apical tips from (B) *bgcn^1/63-44^* versus (C) *bgcn^1/63-44^*; *bam-Gal4 > how RNAi* (VDRC 100775). Arrowhead: spermatocytes. Arrow: spermatid bundles. Scale bars: 50 μm.

**Figure S5. Knockdown of *caf40* in early germ cells by RNAi hairpin resulted in early germ cell overproliferation similar to *bam* mutants.**

(A) Diagram of *Caf40* locus based on Flybase, showing sites of RNAi constructs tested (black lines). (B,C) Phase contrast images of testis apical regions showing two additional examples of *nosGal4 VP16* driving knockdown of *Caf40* using a different *RNAi* line (VDRC 101462). Although the knockdown phenotype was not as strong as for the RNAi construct carried in Bloomington Stock #67987 (Figure 5), there was still overproliferation of small early germ cells followed by cell death. SG: spermatogonia. Arrowhead: spermatocytes. Scale bars: 50 μm. (D,E) Comparison of gene expression levels in testes from *nosGal4;UASCaf40 RNAi* (Bloomington 67987) *versus bam^1/□86^* males 48hrs after heat shock. D) RNA-Seq data showing expression levels per gene plotted as log10(normalized counts + 1). E) Volcano plot comparing RNA-Seq data from (D). (Blue dot) expression of *Caf40*: RNA-Seq confirmed that the knockdown lowered levels of *Caf40* RNA by approximately 14-fold. (Magenta dot) expression of *bam*: The level of *bam* mRNA detected was higher in *Caf40* knockdown than in *bam* mutant testes, likely in part because *bam ^□86^* deletes most of the *bam* coding sequence (26). (Green dot) expression of *how*: RNA-Seq confirmed that levels of *how* RNA in *nosGal4;UASCaf40 RNAi* testes were similar to the levels in *bam* mutant testes.

**Figure S6. Knockdown of *how* in wild-type mid to late spermatogonia did not affect the switch to differentiation.**

(A,B) Phase contrast images of testis apical regions from flies expressing different *how* RNAi hairpins driven by *bamGal4*: (A) Vienna Drosophila Resource Center (VDRC)100775. (B) TRiP HMC03820 from Bloomington Drosophila Stock #55665. Arrowheads: spermatocytes. Arrows: elongating spermatids. Scale bars: 50 μm.

