## Supplemental Information for "An RNA binding regulatory cascade controls the switch from proliferation to differentiation in the *Drosophila* male germ cell lineage"

**This PDF files includes:**

**Figures S1 to S6 and Legends**

**Table 1**

**Table 2**

**Figure S1**

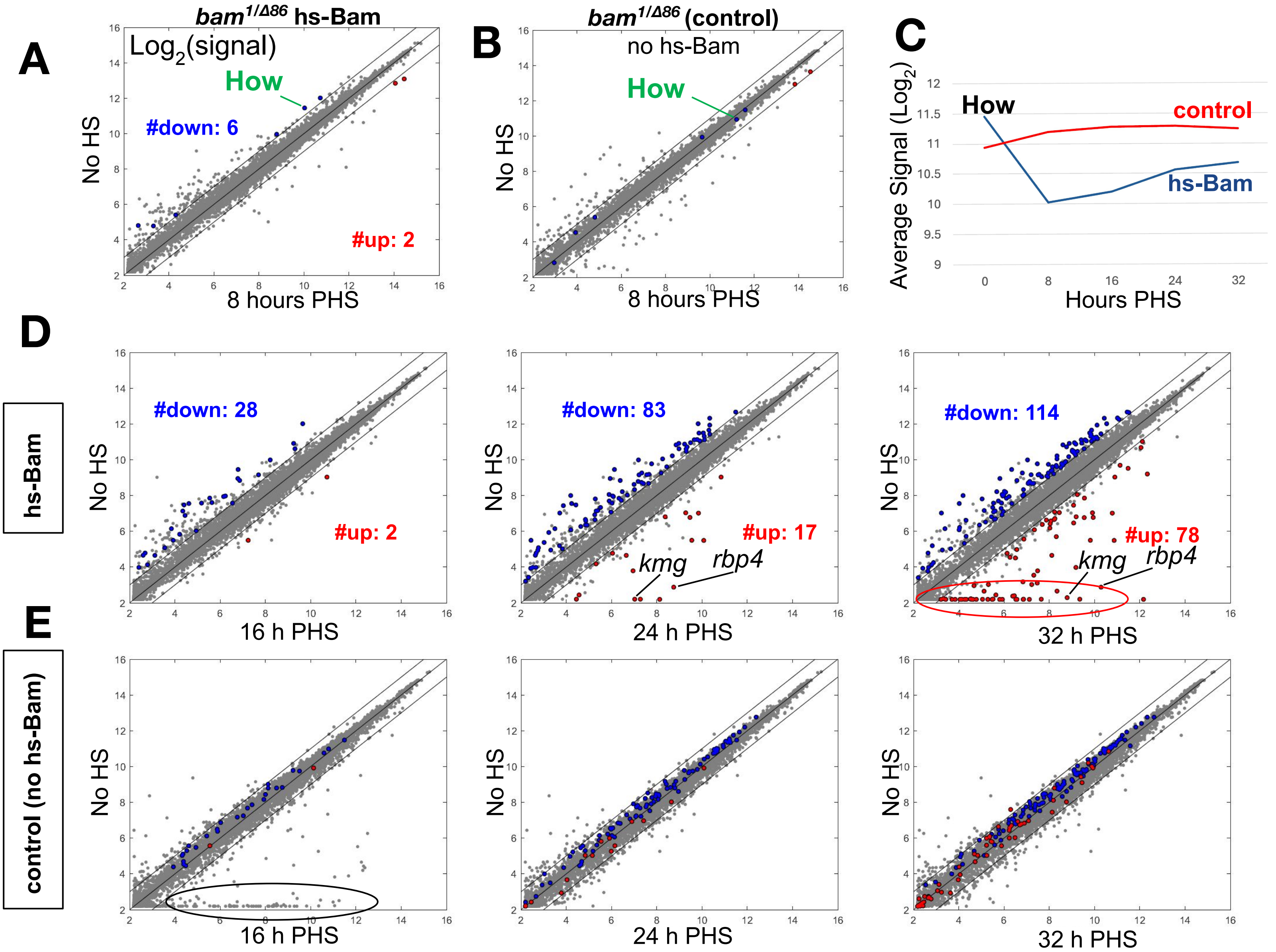
A
B
C
D
E
bam1/Δ86 hs-Bam
Log2(signal)
How
#down: 6
#up: 2
No HS
8 hours PHS
bam1/Δ86 (control)
no hs-Bam
How
No HS
8 hours PHS
How
control
hs-Bam
Average Signal (Log2)
Hours PHS
hs-Bam
#down: 28
#up: 2
16 h PHS
#down: 83
#up: 17
kmg
rbp4
24 h PHS
#down: 114
#up: 78
kmg
rbp4
32 h PHS
control (no hs-Bam)
No HS
16 h PHS
24 h PHS
32 h PHS

**SOM Figure Legends**

**Figure S1.** ***how*** **is among the earliest transcripts to decrease after Bam is turned on.**

(A, B) Scatterplots of transcript levels from microarray analysis comparing no heat shock and 8 hours PHS in (A) *bam*$^{1/\Delta 86}$; *hs-Bam* or (B) *bam*$^{1/\Delta 86}$ males lacking the *hs-Bam* construct, but subjected to the same heat shock regimen in parallel, to control for the effects of heat shock on gene expression. Genes colored blue (downregulated) or red (upregulated) were 1) not changed >2 fold in the microarray analysis of testes from control *bam*$^{1/\Delta 86}$ males lacking the *hs-Bam* construct but heat shocked then incubated for the indicated time and 2) also detected by independent RNA-seq as up or down >2 fold in *bam*$^{1/\Delta 86}$; *hs-Bam* males by RNA-sequencing at the time points indicated compared to testes from flies of the same genotype not subjected to heat shock. (C) Level of *how(L)* transcripts detected by microarray throughout the time course, showing decrease by 8h PHS in flies carrying the *hs-Bam* construct, but not in testes from control *bam*$^{1/\Delta 86}$ flies lacking the hs-Bam construct but subjected to the 30-minute pulse of incubation at high temperature, then shifted back to 25˚C as for the experimental genotype.

(D, E) Microarray data from later time points after heat shock, showing the transcripts that increase in expression (red) and decrease in expression (blue) by the cutoff criteria (detected as up or down regulated >2 fold) in both this microarray comparison and in independent analysis by RNA-seq. (D) *bam*$^{1/\Delta 86}$; *hs-Bam* testes (red oval in *hs-Bam* 32h: genes expressed specifically in spermatocytes. (E) testes from control *bam*$^{1/\Delta 86}$ males that did not have the *hs-Bam* transgene, but were subjected to heat shock then incubated at 25ºC for the indicated times. Black oval in (E) 16h PHS marks genes expressed in accessory glands that contaminated this sample.

#### Figure S2

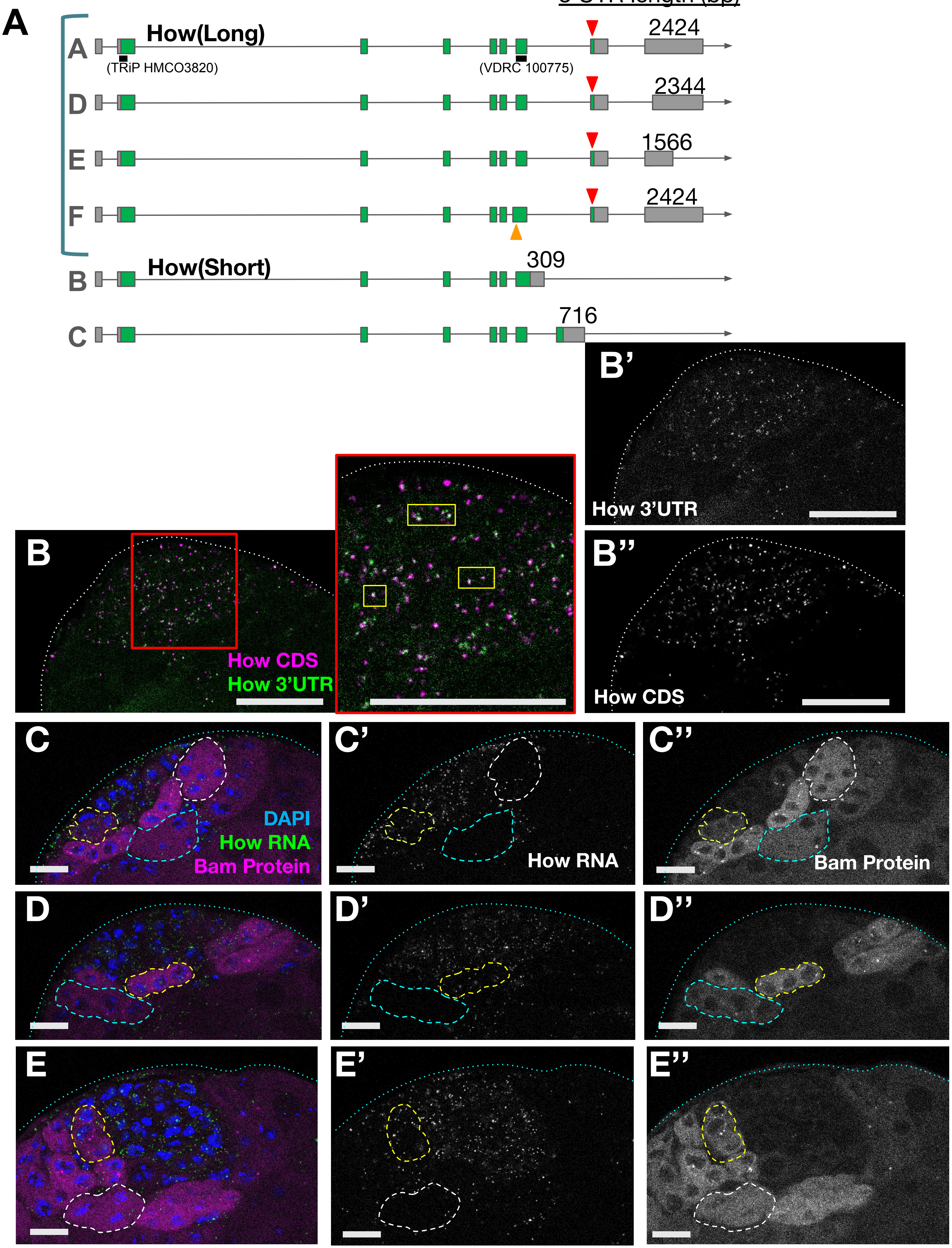

A
3'UTR length (bp)
How(Long)
A
D
E
F
2424
2344
1566
2424
(TRiP HMCO3820)
(VDRC 100775)
How(Short)
B
C
309
716
B
How CDS
How 3'UTR
B'
How 3'UTR
B''
How CDS
C
DAPI
How RNA
Bam Protein
C'
How RNA
C''
Bam Protein
D
D'
D''
E
E'
E''

**Figure S2. How transcripts were detected in spermatogonia but not in spermatocytes**

(A) Diagram of the *how* locus showing mRNA isoforms from FlyBase, designated A-F as in Flybase. The How(L) cDNA construct utilized in Figures 3 and 4 is RA and the How(S) cDNA construct utilized in Figure 3 is RB. Grey: UTRs. Green: protein coding sequence. Lines denote introns. Red arrowheads: nuclear localization signal. Orange arrowhead: additional coding sequence in isoform RF, distinguishing it from RA. Black bars: Sites of RNAi constructs utilized. (B) Apical tip of wild type testis showing distribution of *how* transcripts detected by Hybrid Chain Reaction (HCR FISH) using two different probe sets. (B) Merge. Magenta: Coding sequence (CDS) probe set; Green: 3'UTR probe set. Zoom in marked by red box. Yellow rectangles mark white loci of overlapping probes sets. (B') How(L) 3'UTR probe set. (B") Coding sequence (CDS) probe set. Scale Bar: 25 µm. (C-E) High magnification immunofluorescence images of apical tips of *Bam-GFP* testes with How protein coding sequence RNA labeled by HCR (additional examples for Figure 1G). Left: merge with (blue) DAPI, (green) *how* RNA, and (magenta) Bam-GFP. Middle: *how* RNA only. Right: Bam protein only. Yellow dashed outlines: early Bam positive cysts with *how* RNA present. White dashed outlines: later stage Bam positive cysts with fewer h*ow* RNA foci. Cyan dashed outlines: Bam positive cysts with no or very low *how* RNA signal detected. Scale bar: 12.5 µm.

Figure S3

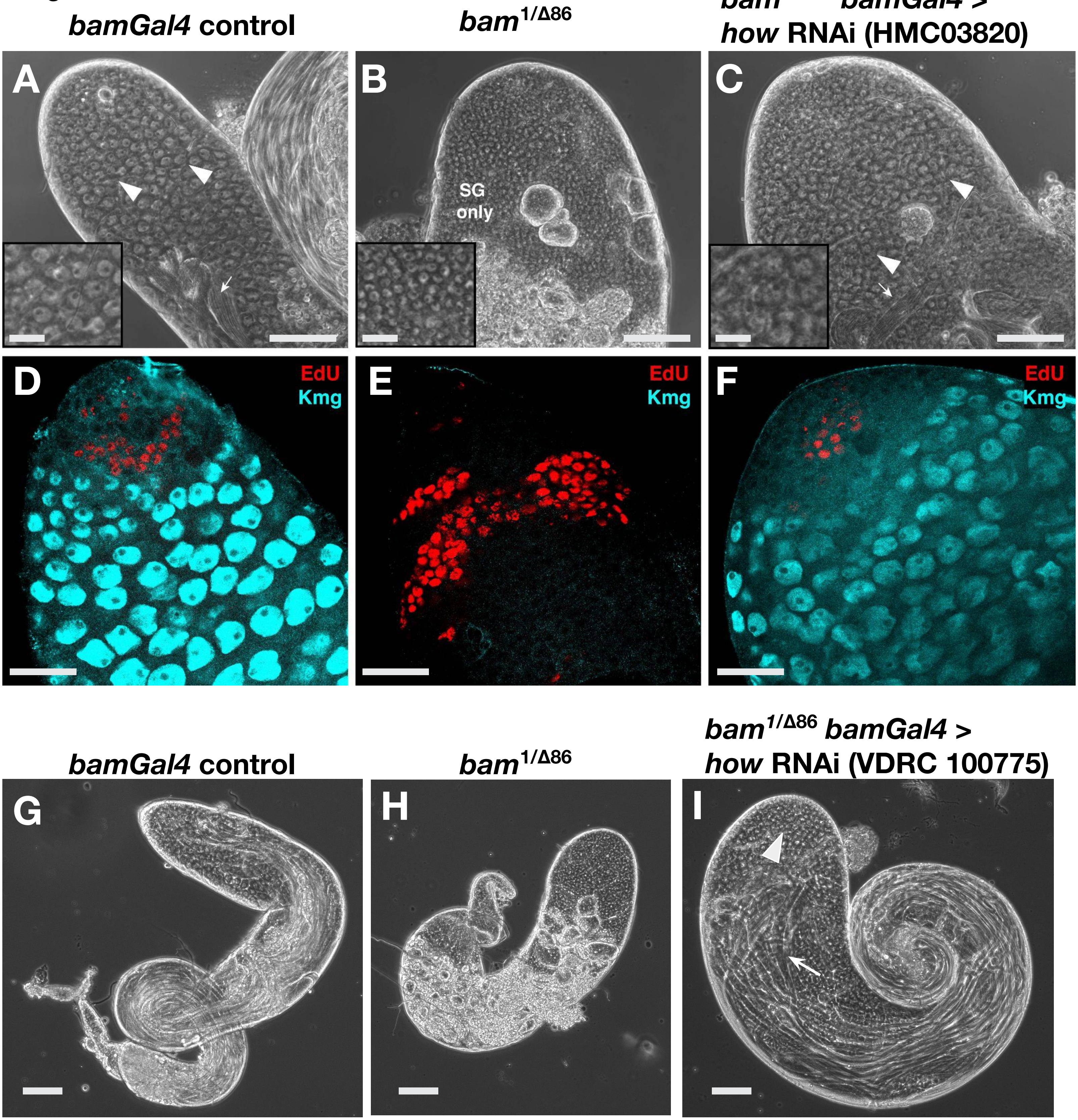


**J**

| Genotype | Fertile Males |
|---|---|
| *bamGal4* | **4/6** |
| *bam*$^{1/\Delta86}$ | **0/3** |
| *bam*$^{1/\Delta86}$ *bamGal4* > *how* RNAi (VDRC 100775) | **5/9** |

**Figure S3. Knockdown of *how* in mid- to late spermatogonia in *bam* mutant males resulted in production of spermatocytes for two different RNAi lines.**

(A-C) Phase contrast images of testis apical tips from (A) *bamGal4* driver only; (B) $bam^{1/\Delta 86}$ ; and (C) $bam^{1/\Delta 86}$; *bamGal4* > *how* RNAi (TRiP HMC03820 from Bloomington Stock #55665). Arrowheads: spermatocytes. Arrows: spermatid bundles. SG: spermatogonia. Scale bars: 100 µm, insets Scale Bars: 25 µm. (D-F) Immunofluorescence images of apical testis tips stained for EdU to mark S-phase nuclei (red) and Kmg to mark spermatocyte nuclei (blue). (D) Control: *bam*Gal4 driver only. (E) $bam^{1/\Delta 86}$. (F) $bam^{1/\Delta 86}$; *bam*Gal4 > *how* RNAi (TRiP HMC03820 from Bloomington Stock #55665). (G-I) Whole testis from (G) *bamGal4* driver only; (H) $bam^{1/\Delta 86}$ ; and (I) $bam^{1/\Delta 86}$; *bamGal4* > *how* RNAi (VDRC100775) for size comparison. Arrowhead: spermatocytes. Arrow: spermatid bundles. Scale bars: 100 µm. (J) Fertility tests of *bamGal4* driver only, $bam^{1/\Delta 86}$; and $bam^{1/\Delta 86}$; *bamGal4* > *how* RNAi (VDRC 100775) males. Males from all genotypes were progeny from crosses where the mated parents were allowed to lay eggs for 3 days at 25˚C then the parents were removed and the progeny shifted to 29˚C. Once the progeny males were collected, they were subjected to fertility tests at RT, ~ 25˚C.

Figure S4

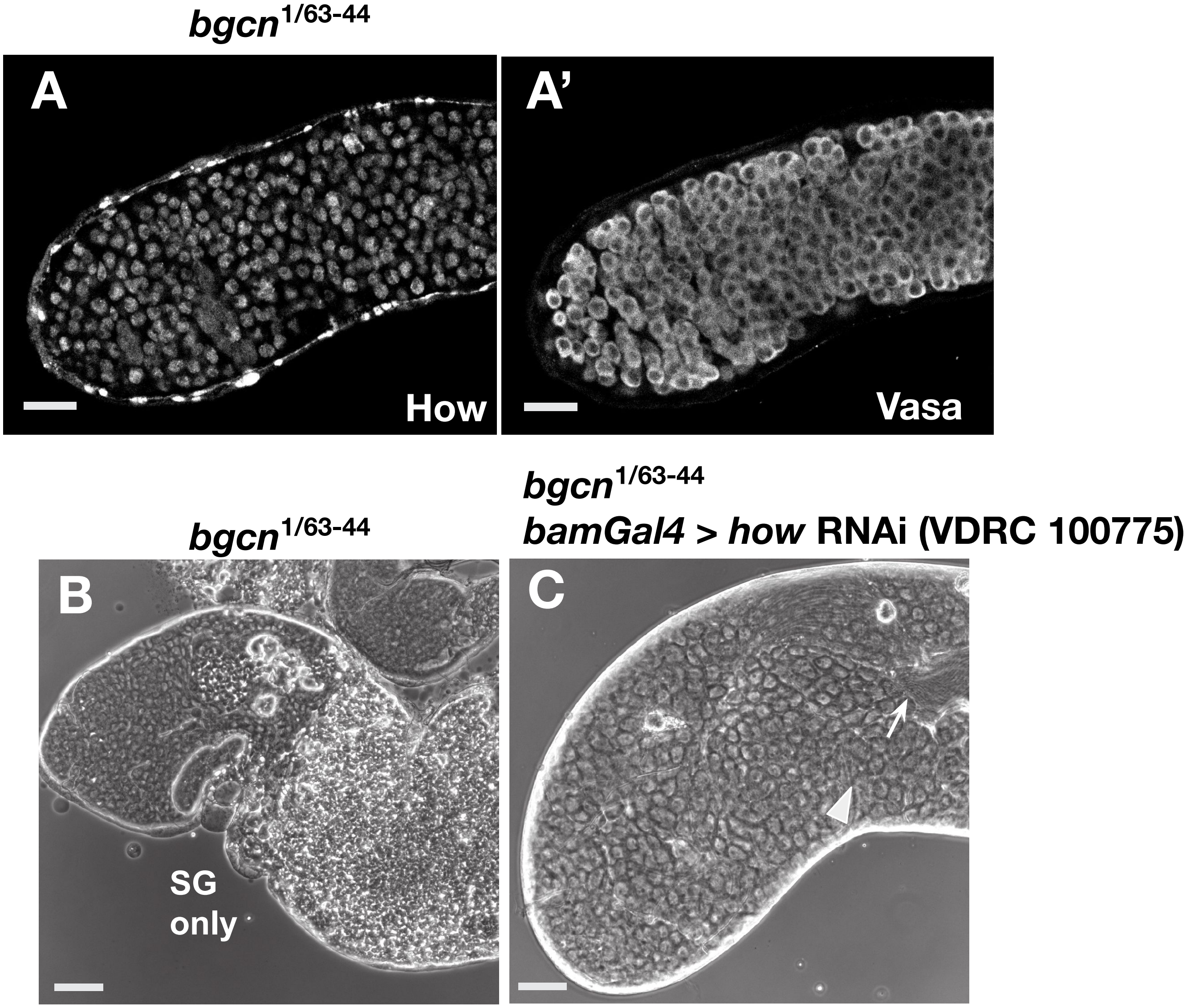

bgcn 1/63-44
A
How
A'
Vasa
bgcn 1/63-44
bamGal4 > how RNAi (VDRC 100775)
bgcn 1/63-44
B
SG only
C

**Figure S4: Knockdown of *how* in mid- to late spermatogonia in *bgcn* mutant males resulted in production of spermatocytes.**

Immunofluorescence images of apical tip of *bgcn*$^{1/63\text{-}44}$ mutant testis stained with (A) anti-How and (A') anti-Vasa. Scale bars: 25 µm. (B,C) Phase contrast images of testis apical tips from (B) *bgcn*$^{1/63\text{-}44}$ versus (C) *bgcn*$^{1/63\text{-}44}$; *bam-Gal4* > *how RNAi* (VDRC 100775). Arrowhead: spermatocytes. Arrow: spermatid bundles. Scale bars: 50 µm.

Figure S5

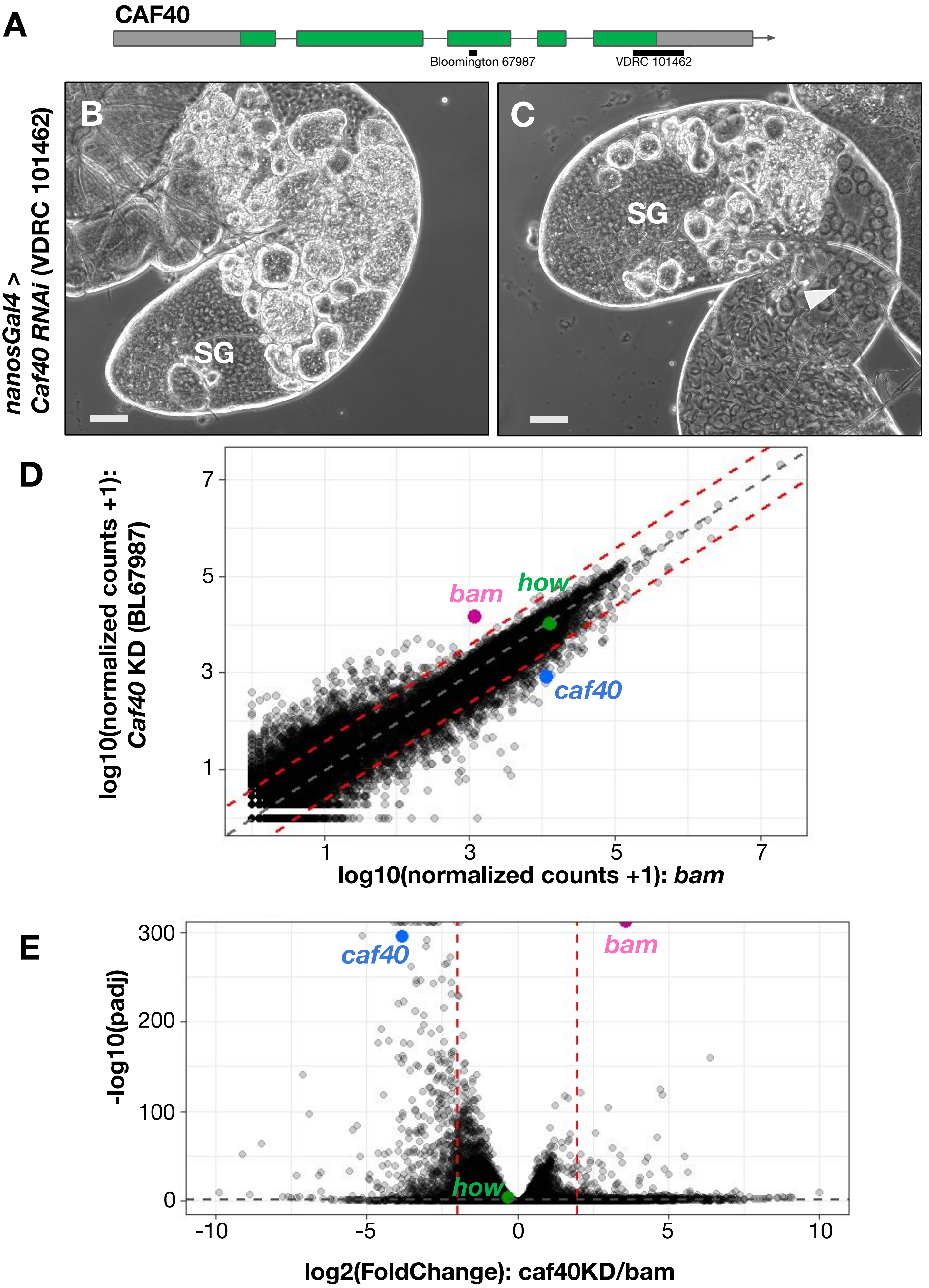
A
CAF40
Bloomington 67987
VDRC 101462
B
C
nanosGal4 >
Caf40 RNAi (VDRC 101462)
SG
SG
D
log10(normalized counts +1):
Caf40 KD (BL67987)
log10(normalized counts +1): bam
bam
how
caf40
1
3
5
7
E
-log10(padj)
log2(FoldChange): caf40KD/bam
caf40
bam
how
300
200
100
0
-10
-5
0
5
10

**Figure S5. Knockdown of *caf40* in early germ cells by RNAi hairpin resulted in early germ cell overproliferation similar to *bam* mutants.**

(A) Diagram of *Caf40* locus based on Flybase, showing sites of RNAi constructs tested (black lines). (B,C) Phase contrast images of testis apical regions showing two additional examples of *nosGal4 VP16* driving knockdown of *Caf40* using a different *RNAi* line (VDRC 101462). Although the knockdown phenotype was not as strong as for the RNAi construct carried in Bloomington Stock #67987 (Figure 5), there was still overproliferation of small early germ cells followed by cell death. SG: spermatogonia. Arrowhead: spermatocytes. Scale bars: 50 µm. (D,E) Comparison of gene expression levels in testes from *nosGal4;UASCaf40 RNAi* (Bloomington 67987) *versus* $bam^{1/\Delta 86}$ males 48hrs after heat shock. D) RNA-Seq data showing expression levels per gene plotted as log10(normalized counts + 1). E) Volcano plot comparing RNA-Seq data from (D). (Blue dot) expression of *Caf40*: RNA-Seq confirmed that the knockdown lowered levels of *Caf40* RNA by approximately 14-fold. (Magenta dot) expression of *bam*: The level of *bam* mRNA detected was higher in *Caf40* knockdown than in *bam* mutant testes, likely in part because $bam^{\Delta 86}$ deletes most of the *bam* coding sequence (26). (Green dot) expression of *how*: RNA-Seq confirmed that levels of *how* RNA in *nosGal4;UASCaf40 RNAi* testes were similar to the levels in *bam* mutant testes.

Figure S6

**A** *bamGal4 > how* RNAi
VDRC 100775

**B** *bamGal4> how* RNAi
TRiP HMC03820
Bloomington 55665

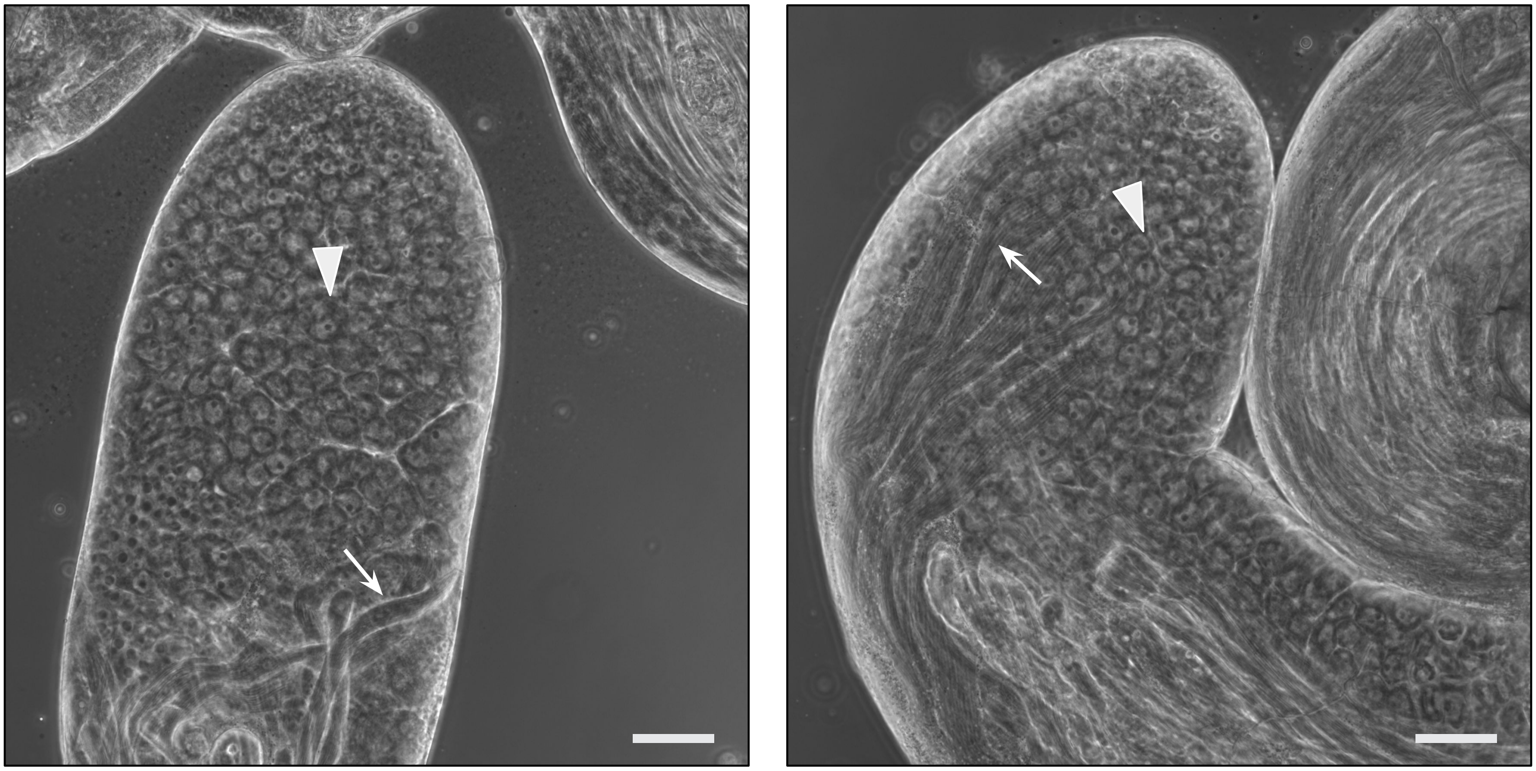

**Figure S6. Knockdown of *how* in wild-type mid to late spermatogonia did not affect the switch to differentiation.**

(A,B) Phase contrast images of testis apical regions from flies expressing different *how* RNAi hairpins driven by *bamGal4*: (A) Vienna Drosophila Resource Center (VDRC)100775. (B) TRiP HMC03820 from Bloomington Drosophila Stock #55665. Arrowheads: spermatocytes. Arrows: elongating spermatids. Scale bars: 50 µm.

### Table 1. *how* coding sequence HCR probes

| |
|---|
| GTCCCTGCCTCTATATCTTTCCTGGGCGGACAATATGCTGGC |
| GGTCGAAGGCGGCTGTCGGAGCTTCCACTCAACTTTAACCCG |
| GTCCCTGCCTCTATATCTTTGGGATTCAGGATCAATGGGGCG |
| CGCCGTTGTGGGGACGGTCATCTTCCACTCAACTTTAACCCG |
| GTCCCTGCCTCTATATCTTTCGGATCTGGGCGGCAAGGCCGG |
| CCAAGCGGGGCGGCGGCGGGTGTTCCACTCAACTTTAACCCG |
| GTCCCTGCCTCTATATCTTTGGCTATCAGAGGCGGCAACCAG |
| GCAGGCCGGTGGATGTCAGCAGTTCCACTCAACTTTAACCCG |
| GTCCCTGCCTCTATATCTTTTGCGACAGATTTCGCTGTTGTG |
| GCGGCGCCACTCCTCATCGCACTTCCACTCAACTTTAACCCG |
| GTCCCTGCCTCTATATCTTTGCCAATTCCATGAGTTGACGTT |
| TCCCTATAAGTGCCATTAATAATTCCACTCAACTTTAACCCG |
| GTCCCTGCCTCTATATCTTTCTTGCGGCACGAGCAACTTCTG |
| TCTTTAGCTCATCTTCGCCTTCTTCCACTCAACTTTAACCCG |
| GTCCCTGCCTCTATATCTTTCTTCACTGTGGCACGGTTCTCG |
| TACTTCGGCGACGGCCTGGGCCTTCCACTCAACTTTAACCCG |
| GTCCCTGCCTCTATATCTTTTGCAGGTCATCGGACAGATGCT |
| GTGTCCTCGACGGTTATCAGGATTCCACTCAACTTTAACCCG |
| GTCCCTGCCTCTATATCTTTCGTCCTCCTTCTTCTTGTCGCG |
| CCCAGTTAGGCTTGCCACGGTTTTCCACTCAACTTTAACCCG |
| GTCCCTGCCTCTATATCTTTAATCTTGCAGCCGGTCTCCTGT |
| CATGGAACCCTTGCCTCGGACCTTCCACTCAACTTTAACCCG |
| GTCCCTGCCTCTATATCTTTCGGGGTCCCAAAATGCGACCGA |
| TCCAATTGCTTGGCGGTCATGCTTCCACTCAACTTTAACCCG |
| GTCCCTGCCTCTATATCTTTGGACTGGCACATAAACCTTCTC |
| CAAAGTTGAAATCTGGATGCTCTTCCACTCAACTTTAACCCG |
| GTCCCTGCCTCTATATCTTTGGGTTCGGGCAGAGTGAGCGGC |
| GTTCATCGTCACCACAGAGCCCTTCCACTCAACTTTAACCCG |
| GTCCCTGCCTCTATATCTTTAACAGTGAGGCGCGCACGCGTG |
| TCCTTCTTGACCCCATTGATCTTTCCACTCAACTTTAACCCG |
| GTCCCTGCCTCTATATCTTTCGACGTGGGTGAAGACGTTGGG |
| CAATTTCTTCGTCCAGCAGGCGTTCCACTCAACTTTAACCCG |
| GTCCCTGCCTCTATATCTTTCTTGAGCAACTGGGCCAGATAG |

#### Table 1. continued

| |
|---|
| GAAGGCGGCCAGTTGCTTGCGGTTCCACTCAACTTTAACCCG |
| GTCCCTGCCTCTATATCTTTTGCTGCTGCTGTGGGGTCAAGT |
| TCGGCGATGCTCTGTGTGCTCTTTCCACTCAACTTTAACCCG |
| GTCCCTGCCTCTATATCTTTCCTGCGGCGCCTGTTGCTGCTG |
| GCTGCGGGGTCATGGGGACCACTTCCACTCAACTTTAACCCG |
| GTCCCTGCCTCTATATCTTTTTGGGCTTGAGCCTGCTGCTGT |
| CTGTGCCTGAGCCTGAGCCTGGTTCCACTCAACTTTAACCCG |
| GTCCCTGCCTCTATATCTTTGCTGCCTGTTGCTGCAAGTGCT |
| TGCGCGACCGCAACAACTGCTGTTCCACTCAACTTTAACCCG |
| GTCCCTGCCTCTATATCTTTCTTTGCTCTCACAGACACTCAT |
| GCTGCAGTTGCTGTTGCACAACTTCCACTCAACTTTAACCCG |

### Table 2. *how(L)* 3’UTR HCR probes

| |
|---|
| CTCACTCCCAATCTCTATAATATATTTAAATATATATATATA |
| CTTGTATATGCATTTGGATGTAAACTACCCTACAAATCCAAT |
| CTCACTCCCAATCTCTATAATTTTGTTCGTTGAACTTTTACA |
| TGTATATGCTTGCATTAAGTATAACTACCCTACAAATCCAAT |
| CTCACTCCCAATCTCTATAATTAGATTATCATAATTAAACTA |
| CTAAGAACTAACTTGTATTTTCAACTACCCTACAAATCCAAT |
| CTCACTCCCAATCTCTATAAAATCGTTTGCTTAATTGCCTTG |
| AGCTCTATGTTTGATTCTCTTAAACTACCCTACAAATCCAAT |
| CTCACTCCCAATCTCTATAAAGTTTAATGGCCACAAGATTCA |
| TTTTGTTTGCTGGCAAAACGTAAACTACCCTACAAATCCAAT |
| CTCACTCCCAATCTCTATAACGTTTGCTGTTGCTGTTGCTTT |
| ATACATACTGATGTTGCTGCTGAACTACCCTACAAATCCAAT |
| CTCACTCCCAATCTCTATAATGTGATGCTGCTGGTGTTGAAT |
| TGCTTTTGCTTTTGCTGTTGATAACTACCCTACAAATCCAAT |
| CTCACTCCCAATCTCTATAACAAATAGGTTCAGAGTTTGGAG |
| GTGCCGCATCTGCTGCTCATTTAACTACCCTACAAATCCAAT |
| CTCACTCCCAATCTCTATAATTTTAGTAGTTGTGTAAGTGAC |
| TTTGGCGTTCGGAGGGGTTCAGAACTACCCTACAAATCCAAT |
| CTCACTCCCAATCTCTATAATTTTGGTTACGCTTTTTTGTTA |
| ATGTGGTGGCTCAGGATATCTGAACTACCCTACAAATCCAAT |
| CTCACTCCCAATCTCTATAAACACACGTGGCTCTGCTTTCGG |
| GCTTCAAGTAAGCACATTCACAAACTACCCTACAAATCCAAT |
| CTCACTCCCAATCTCTATAACTGATTCCGTGCTCATCTACGG |
| CCTGGCTGGGATGCTGCTGCTGAACTACCCTACAAATCCAAT |
| CTCACTCCCAATCTCTATAAGCGGATGTGGGAGCGGATTGGT |
| CAGGCGATGGATTCATGGTTCAAACTACCCTACAAATCCAAT |
| CTCACTCCCAATCTCTATAATTTCCTGTGGAGGCGGTGCGCT |
| AGTGTGTTTGGGACATTTGTTTAACTACCCTACAAATCCAAT |
| CTCACTCCCAATCTCTATAAATCAGGGCAGAGCGGCCAGGCC |
| TGAAAATTCTTTTTCAGAACCCAACTACCCTACAAATCCAAT |
| CTCACTCCCAATCTCTATAACATCGAACATACAGTGTGGATT |
| AGGCCAACGATTCCGGTCAGGTAACTACCCTACAAATCCAAT |
| CTCACTCCCAATCTCTATAAGCAGCATTTGTATCAAAATAAA |
| GGAATTGGGGATGGTCAGTTATAACTACCCTACAAATCCAAT |
| CTCACTCCCAATCTCTATAACTAAAGTGATCGGCTGCCTGGC |

### Table 2. continued

| |
|---|
| TAGCAAAATTTTAAAATAATTGAACTACCCTACAAATCCAAT |
| CTCACTCCCAATCTCTATAACGCAATCGTTCGTTATCCGTTG |
| GTACGCAATCAAAGGGCAGGCAAACTACCCTACAAATCCAAT |
| CTCACTCCCAATCTCTATAATCGTACATTATTGATTTAAAGT |
| TTCTTAATGGTTTCTTTTCTTCAACTACCCTACAAATCCAAT |
| CTCACTCCCAATCTCTATAAGTGTGATTTTAGAGTGTTACTT |
| TATAAAGTTGCTTTCGCCGATAAACTACCCTACAAATCCAAT |
| CTCACTCCCAATCTCTATAACTCGATTCACTATGCATGGTAT |
| TCTGAGTTATAGTGCTTCGAATAACTACCCTACAAATCCAAT |
